# Xenometabolome of Early-Life Stage Salmonids Exposed to 6PPD-Quinone

**DOI:** 10.1101/2024.06.12.598661

**Authors:** Phillip J. Ankley, Francisco C. da Silva, Blake Hunnie, Catherine Roberts, Andreas N. M. Eriksson, Evan Kohlman, Jennifer Corker, Jodi Westcott, Bradlee Germain, Michael J. Martin, Justin Dubiel, Katherine Anderson-Bain, Rayen M. Urrutia, Natacha Hogan, John P. Giesy, Steve Wiseman, Ed Krol, Markus Hecker, Markus Brinkmann

## Abstract

*N*-(1,3-Dimethylbutyl)-*N′*-phenyl-*p*-phenylenediamine-quinone (6PPD-Q) is a ubiquitous transformation product (TP) derived from the rubber tire antioxidant *N*-(1,3-Dimethylbutyl)-*N′*-phenyl-*p*-phenylenediamine (6PPD) and is acutely toxic to certain species of *Salmonidae*. Not all salmonids are sensitive to acute lethality caused by 6PPD-Q, with 6PPD-Q potency varying by several orders of magnitude among teleosts. The main driver(s) of species sensitivity differences is (are) a pressing question, with one area of interest examining whether differences in teleosts ability to biotransform and detoxify 6PPD-Q could be a key factor. This study utilized liquid-chromatography high-resolution mass spectrometry (LC-HRMS) to assess biotransformation and metabolome-wide effects of 6PPD-Q on early-life stage salmonids, including two sensitive species, rainbow trout (*Oncorhynchus mykiss*) and lake trout (*Salvelinus namaycush*), and one tolerant species, brown trout (*Salmo trutta*). Three phase I TPs and seven phase II TPs were detected, with differences in peak areas revealing that brown trout had the greatest ability to detoxify 6PPD-Q. Monohydroxylated TPs were verified using co-developed analytical standards that will be of use for future biomonitoring and exposure assessment. Several endogenous metabolites were found to be dysregulated in rainbow and lake trout, indicative of mitochondrial dysfunction, altered metabolism, and disrupted membrane permeability. Results of this study indicate a difference in the biotransformation capability of 6PPD-Q among *Salmonidae* fish species and subsequent unique metabolome responses.

**Graphical Abstract:** 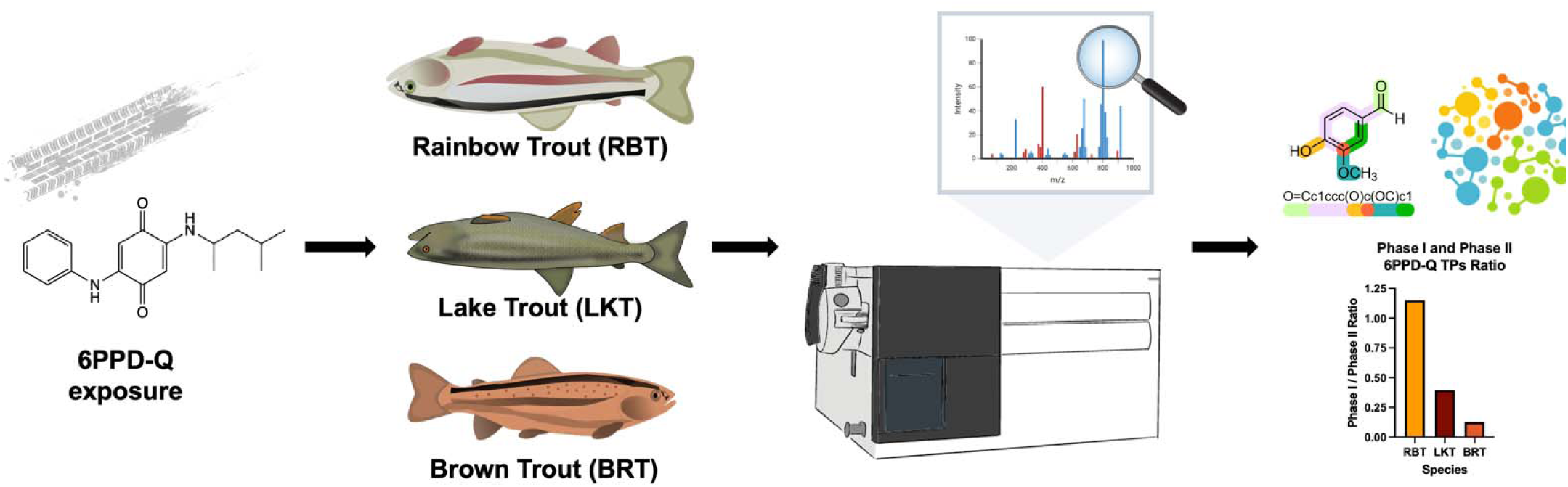

**Synopsis:** Inter-species salmonid differences in the biotransformation capability of 6PPD-Q leads to sensitivity variability and subsequent unique metabolome responses.

## INTRODUCTION

*N*-(1,3-Dimethylbutyl)-*N′*-phenyl-*p*-phenylenediamine-quinone (6PPD-Q) is a ubiquitous and *Salmonidae*-specific acutely toxic transformation product (TP) derived from the rubber tire antioxidant *N*-(1,3-Dimethylbutyl)-*N′*-phenyl-*p*-phenylenediamine (6PPD).^1^ 6PPD is released into the environment through diffusion from and wear of tire surfaces, where it can react with oxidants, such as ozone, to form 6PPD-Q.^2^ Urban runoff after rainfall events is the main source of 6PPD-Q into receiving aquatic ecosystems.^3^ The total global amount of 6PPD-Q generated per year is estimated to fall between 26 to 1,900 tons,^4^ with this chemical being detected globally up to levels of 2.29 µg/L in aquatic environments.^1^ The amount of this contaminant being released into aquatic environments, and concentrations at which it has been detected *in situ*, have been shown to impact fish populations adversely. In particular, the coho salmon (*Oncorhynchus kisutch*), and other select *Salmonidae*, have been shown to be especially sensitive to 6PPD-Q exposure.^1,5^

6PPD-Q is the leading cause of urban runoff mortality syndrome (URMS) in U.S. Pacific Northwest coho salmon,^1^ with a 24-h median lethal concentration (LC_50_) of 0.095 µg/L.^6^ 6PPD-Q has also been shown to be acutely toxic to other salmonids, including adult brook trout (*Salvelinus fontinalis*; 24-h LC_50_ = 0.59 µg/L),^6^ rainbow trout (*Oncorhynchus mykiss*; 24-h LC_50_ = 1.96 µg/L),^5^ white-spotted char (*Salvelinus leucomaenis pluvius*; 24-h LC_50_ = 0.8 µg/L),^7^ and juvenile lake trout (*Salvelinus namaycush*; 96-h LC_50_ = 0.50 µg/L).^8^ In contrast, most other fish species tested to date, such as white sturgeon (*Acipenser transmontanus*; 24-h LC_50_ > 12.7 µg/L)^5^ and zebrafish (*Danio rerio*; 24-h LC_50_ = 308.7 µg/L),^9^ as well as other salmonids including arctic char (*Salvelinus alpinus*; LC_50_ > 14.2 µg/L)^5^ and brown trout (*Salmo trutta*; 48-h alevin LC_50_ > 12.2 µg/L),^10^ are several orders of magnitude less sensitive to aqueous exposure of 6PPD-Q. The specific factors driving these large species differences in sensitivity are a pressing question for ecotoxicologists and risk assessors.

One area of investigation regarding a potential determinant of differences in sensitivity among fish species is their ability to biotransform and detoxify 6PPD-Q upon exposure. Previous research has indicated that fish differ in the levels of TPs from 6PPD-Q exposure, including increased levels of phase I hydroxylated 6PPD-Q and phase II glucuronidated 6PPD-Q in the bile of more tolerant species, i.e., chinook salmon (*Oncorhynchus tshawytscha*), westslope cutthroat trout (*Oncorhynchus clarkii lewisi*), and white sturgeon.^11^ Zebrafish embryos have also been shown to have multiple biotransformation pathways, including sulfonate-, N-acetylcysteine-, and glutathione-conjugation, that could reduce sensitivity to 6PPD-Q by effectively detoxifying the chemical.^12^ Hydroxylated 6PPD-Q TPs are a result of phase I biotransformation, including an alkyl sidechain monohydroxylated TP (TP-OH1), a phenyl ring monohydroxylated TP (TP-OH2),^13^ and a dihydroxylated TP (TP-2-OH).^13,14^ However, little is known about additional phase II TPs occurring in 6PPD-Q-exposed salmonids, which should be better characterized to understand differences in detoxification upon aqueous exposure.

Non-targeted liquid-chromatography high-resolution mass spectrometry (LC-HRMS) provides a powerful method to characterize the xenometabolome and endometabolome of tissue of larval fishes.^15,16^ Xenometabolome analysis profiles TPs from the biotransformation of contaminants within organisms from environmental exposure,^17^ allowing the understanding of exposure and metabolism of contaminants that could lead to potential toxicity (reactive metabolite) or detoxification (inactive metabolite). Endogenous metabolites are involved in the growth and homeostasis of fish, and the metabolome profile is the ultimate measurement of cellular and tissue responses to toxicant exposure and a key link associating genes with observable phenotypes.^18,19^ Non-targeted LC-HRMS has the capability to simultaneously profile both the xenometabolome and the endometabolome of fishes for assessment of contaminant contribution to toxicity and changes in fish health from exposure.^20–22^

This study utilized LC-HRMS to assess biotransformation and metabolome-wide effects of 6PPD-Q on the early-life stages of three salmonids, including two sensitive species, rainbow trout (28-d alevin LC_50_ = 0.56 µg/L)^23^ and lake trout (45-d alevin LC_50_ = 0.39 µg/L),^8^ and one tolerant species, brown trout.^23^ Specific objectives of this study included: (1) profile the xenometabolome of 6PPD-Q within the three selected salmonids; (2) compare and verify detected TPs and levels among the species; and (3) assess if and how 6PPD-Q affects the endometabolome of each species.

## MATERIALS AND METHODS

### Chemicals and Fish Culture

6PPD-Q (CAS #: 2754428-18-5) and deuterium-labeled 6PPD-Q-d5 (CAS #: 2750119-14-1) were purchased from Toronto Research Chemicals (Canada) and had purities of ≥ 97%, with standard solutions being prepared in HPLC-grade methanol (MeOH). Stock solutions for fish exposures were prepared in carbon- and bio-filtered City of Saskatoon water and dimethyl sulfoxide (DMSO) to achieve a final solvent concentration of 0.1% (v/v).

Lake trout (LKT) embryos were obtained following field spawning from White Swan Lake, Saskatchewan, and transported to the Aquatic Toxicology Research Facility (ATRF) at the University of Saskatchewan (SK, Canada) under the Government of Saskatchewan Fish Transport/Stocking permit FTP02-2022, as described previously.^24^ Eyed embryos were moved into glass aquaria, held under a 12:12 h light: dark photoperiod, and gradually acclimated to 10 ± 0.5 °C until hatch. Triploid rainbow trout (RBT) eyed embryos were obtained from Lyndon Fish Hatcheries (ON, Canada). Eyed embryos were kept in an environmentally controlled exposure chamber in glass aquaria with aerated dechlorinated facility water and acclimated as described above using temperatures of 14 ± 0.5 °C until hatch. Brown trout (BRT) embryos were obtained from the Allison Creek Brood Trout Station (Coleman) in Alberta, Canada. Approximately 4000 newly fertilized embryos were transported to the Aquatic Research Facility (ARF) at the University of Lethbridge (AB, Canada) under dark and cool conditions and maintained in a flow-through system at a temperature of 10 °C and 16:8 h light: dark photoperiod until hatching.

### Exposure Experiments

All fish experiments were conducted independently, and varying nominal concentrations were used for exposure (Supplementary Table S1). LKT and RBT studies were approved by the University of Saskatchewan Animal Research Ethics Board (Animal Use Protocol 2022-0002) following Canadian Council on Animal Care guidelines and greater details of the experiments can be found in previous publications.^8,23^ BRT exposures were approved by the University of Lethbridge Animal Welfare Committee (Animal Use Protocol 2324).

Newly hatched LKT, RBT, and BRT were exposed to nominal concentrations (ranging from 0.125 µg/L – 10 µg/L) of 6PPD-Q (Supplementary Table S1) until two weeks post-swim up following OECD guidelines for the Early-life Stage Toxicity Test No. 210.^24^ Upon hatch, RBT and LKT had 18 and 15 alevins, respectively, placed in each tank (2.5 L) with five replicate tanks for each concentration (RBT – 0.125, 0.25, 0.5, 1, 2, and 4 µg/L; LKT – 1.25 and 2.5 µg/L) as well as a solvent control of 0.01% DMSO. BRT had 30 alevins in each tank (3.5 L) with five replicate tanks per chemical treatment (2.5, 5, and 10 µg/L). Fish were maintained at an average daily temperature of 10 °C for LKT, 14 °C for RBT, and 10 °C for BRT, with a photoperiod of 12:12 h light: dark, gaining 1 hr light per week for a final photoperiod of 16:8 h light: dark for LKT, RBT, and BRT All exposures were conducted under semistatic-renewal conditions, with 70% replacement of water and chemicals daily throughout the entire exposure. Water quality parameters were monitored daily in randomly selected aquaria, including temperature and dissolved oxygen, and incoming processed facility water for ammonia, nitrates, and nitrites (Supplementary Table S2).^8^ Samples of water were collected into LC vials throughout the exposure and stored at –20 °C to confirm concentrations of chemicals using LC-HRMS as described previously,^8^ spiking 950 µL of water with 50 µL of 1000 µg/L 6PPD-Q-d5 for a final internal standard concentration of 50 µg/L (Supplementary Text S1). Salmonids were fed with hatched brine shrimp (*Artemia* spp. nauplii) once daily approaching and through the first week of swim-up and twice daily for the remainder of the exposure until 24 hours prior to termination. At 28 days post-hatch (dph) for RBT and BRT and 45 dph for LKT, fry were euthanized in 150 mg/L buffered tricaine methanesulfonate (MS-222). Individual fish were weighed and measured for length (standard and total). Two to three fish from each tank were randomly collected for xenometabolome analysis of whole larval tissue.

### Mass Spectrometry Analysis

Whole larval fish were weighed before and after freeze-drying, a process that lasted for 48 h. Twelve volumes of MeOH (µL) per mg of dried mass (dm) were added with the inclusion of two 5-mm steel beads and 25 µL of 6PPD-Q-d5 dissolved in MeOH (1,000 µg/L) before complete homogenization using Tissue Lyzer II at 20 Hz for 3 minutes. Samples were then centrifuged at 4,000 × g at 4 °C for 30 min with the supernatant transferred to a new 2.0 mL microcentrifuge tube with 6 volumes of ACN and vortexed for 1 min before storage at –20 °C for 12 h. After overnight salt and protein precipitation, samples were centrifuged at 4,000 × g at 4 °C for 30 min before transferring the supernatant to a new 2.0-mL glass vial to undergo nitrogen drying. Samples were then reconstituted in 500 µL of 50:50 MeOH: H_2_O and filtered using a syringe + PTFE 0.22 µM syringe filter (Agilent, USA) into a new 2-mL glass vial before LC-HRMS analysis with a final internal standard concentration of 50 µg 6PPD-Q-d5/L.

Non-targeted analyses were conducted utilizing a data-dependent acquisition (DDA) method. Details on the separation of analytes using ultra-high performance liquid chromatography (UHPLC) coupled with a heated electrospray ionization (HESI) ion source to a Q-Exactive^TM^ HF Quadrupole-Orbitrap (Thermo Scientific) are provided in Supplementary Text S3 and Table S4. The following parameters for non-targeted analysis of extracts of larval fish were used to acquire the ddMS2 top 5 positive mode scans: sheath gas flow = 35, aux gas flow = 10, sweep gas flow = 1, aux gas heater temperature = 300 °C, spray voltage = 3.00 kV, S-lens RF = 60.0, and capillary temperature = 350 °C. A Full MS method used scan settings of resolution = 60,000, positive ion mode, AGC target = 5e5, maximum injection time = 100 ms, and Full MS scan range of 70-1,000 *m/z*. The ddMS2 used the following scan settings: resolution = 15,000, AGC target = 1e5, maximum injection time = 100 ms, loop count = 5, TopN = 5, an isolation window of 2.0 *m/z*, and stepped normalized collision energy (NCE) of 15, 30 and 45. Blanks and a quality control (QC) pool of all samples for each species were run throughout the sequence, and a 4-point standard curve was run at the beginning of each run (0.05, 2.5, 5, and 25 µg/L) for semi-quantification of 6PPD-Q body burdens. Monohydroxylated TPs were confirmed and distinguished using co-developed analytical standards from Cayman Chemical using a paired full scan and parallel reaction monitoring (PRM) method using the same LC-HRMS setup (Supplementary Text S3 and S4). These synthesized analytical standards were compared to biologically occurring monohydroxylated TPs for verification (Supplementary Text S3).

### Non-targeted Data Analysis

Thermo *.raw files were converted to mzML format using MSConvert with the peak picking filter before downstream processing using MZmine 3.^25^ Spectral preprocessing, feature detection, and preliminary compound identification using MoNA LC-MS-MS Positive Mode database (Accessed August 23, 2023) were conducted in MZmine 3 (version 4.0.3) using parameters outlined in supporting information (Supplementary File S1). To eliminate background, features were removed if detected in two or more blanks and were less than 10-fold more abundant in samples. After processing, mgf files were exported using the export feature lists command for feature-based molecular networking (FBMN) using GNPS web server and molecular formula identification and compound annotation using SIRIUS (version 5.8.5).^26,27^ FBMN was run using default settings, and the resulting graph was exported and visualized in Cytoscape to detect and prioritize TPs.^28^ Several new TPs were observed and identified using Biotransformer 3.0 and CFM-ID for spectral verification, along with review of literature.^13,29^ SIRIUS GUI was used to analyze the feature data using the SIRIUS algorithm with Orbitrap-based default settings. Features were additionally analyzed using CSI:FingerID with Fingerprint Prediction using Fallback Adducts of [M+H]+, [M]+, [M+K]+, and [M+Na]+, and Structure Database Search using Bio Database, HMDB, and KEGG.^30^

The resulting data was filtered and merged to detect TPs and compare differences among species for the 2-2.5 µg/L 6PPD-Q exposure bin. TPs were kept if they were detected in QC samples, omitting TPs detected at low abundance and in a small number of samples. TPs that shared identical *m/z* were merged if retention times (RTs) were ± 0.5 min of the average and had the same fragmentation pattern of precursor, indicating they were the same TP with a slight shift in RT, which can occur at greater concentrations when conducting a DDA experiment and when running multiple experiments. Details of precursor and product ions used for verification are provided in the supplementary information (Supplementary Table S5). If merged, the peak area was summed, and the RT was averaged for the respective TPs. Peak areas were normalized to measured dm of larval fish to compute the normalized relative abundance of TPs, as fish species differed in size upon take-down (Supplementary Table S1). To investigate the relative bioaccumulation between salmonid species, the 4-point standard curve was used to estimate 6PPD-Q body burden from peak areas, with concentrations being normalized to dm of fish.

### Statistics

Data analyses and visualizations of 6PPD-Q body burden and normalized peak areas of TPs were performed using GraphPad Prism software (version 10.1.1). MetaboAnalyst (version 6.0) was used to analyze larval fish metabolome response to 6PPD-Q.^31^ All features passing filtering were used to compute a Partial Least-Squares Discriminant Analysis (PLS-DA) to discern potential differences between treatments (Supplementary Text S5). A volcano plot was used to differentiate dysregulated metabolites between solvent control and 2-2.5 µg/L exposure bin using a false-discovery rate (FDR) *p*-value of 0.05 and fold change (FC) of 2.0. Significant features that could be annotated by having a compound name derived from SIRIUS and mapped to MetaboAnalyst common metabolite name database were used to further interpret the biological response (Supplementary File S2). Concentration-response analysis computed metabolite benchmark doses (BMDs) for metabolomic points-of-departure (mPOD) using default settings while including Exp5 and Hill statistical models with Exp2, Exp3, Exp4, Linear, Poly2, and Power models.^31^ Briefly, an ANOVA was performed using a limma R package, with metabolites having an FDR < 0.05 fitted to statistical models. BMDs were calculated using a lack-of-fit p-value of 0.10, benchmark response factor of 1.0, and control abundance using the mean of control samples. mPODs were estimated from the metabolomic BMD analysis as: (1) the BMD of the 20^th^ most sensitive metabolite, (2) the max 1st peak of metabolite BMD distribution, and (3) the 10^th^ percentile of all metabolite BMDs.

## RESULTS AND DISCUSSION

### Levels of 6PPD-Q Within Fish Samples

Verification of nominal exposure concentrations for 2-2.5 µg/L exposure bin indicated time-weighted averaged concentrations of 1.30 ± 0.250 µg/L (average ± standard deviation) (65.0% of nominal exposure concentration),^23^ 1.30 ± 0.167 µg/L (52.0%),^8^ and 0.615 ± 0.550 µg/L (24.6%) for RBT, LKT, and BRT, respectively. Recovery of 6PPD-Q-d5 from larval fish extracts was 37.0 ± 7.84%, 68.9 ± 18.4%, and 47.1 ± 8.11%. for RBT, LKT, and BRT, respectively. Methanol was used to prioritize polar metabolites and TPs; therefore, some loss of 6PPD-Q and 6PPD-Q-d5 (RT = 15.7) was expected.^32^ 6PPD-Q body burdens were 33.2 ± 24.1 ng/g (RBT), 41.7 ± 13.0 ng/g (LKT), and 44.9 ± 25.5 ng/g (BRT) for the 2-2.5 µg/L exposure bin. Concentrations of 6PPD-Q body burdens increased among fish species with increasing exposure concentrations (Figure 1A). At the 2-2.5 µg/L exposure, LKT and BRT tended towards slightly greater body burdens of 6PPD-Q compared to RBT (Figure 1B; ANOVA, *p*-value = 0.746), but these likely resulted from the difference in nominal exposure concentrations (2 µg/L – RBT vs. 2.5 µg/L – LKT and BRT). More research that incorporates time-course sampling of larval fish from aqueous exposure is needed to understand the rate and magnitude of 6PPD-Q bioaccumulation between species. Bioaccumulation is an important measurement for environmental risk assessment,^33^ and exposure-time is a necessary endpoint to consider when assessing bioaccumulation of chemicals. Additionally, 6PPD-Q is known to have time-driven effects,^1,34^ with research suggesting differences between acute and sub-chronic mode of toxic action.^8^

**Figure 1.**
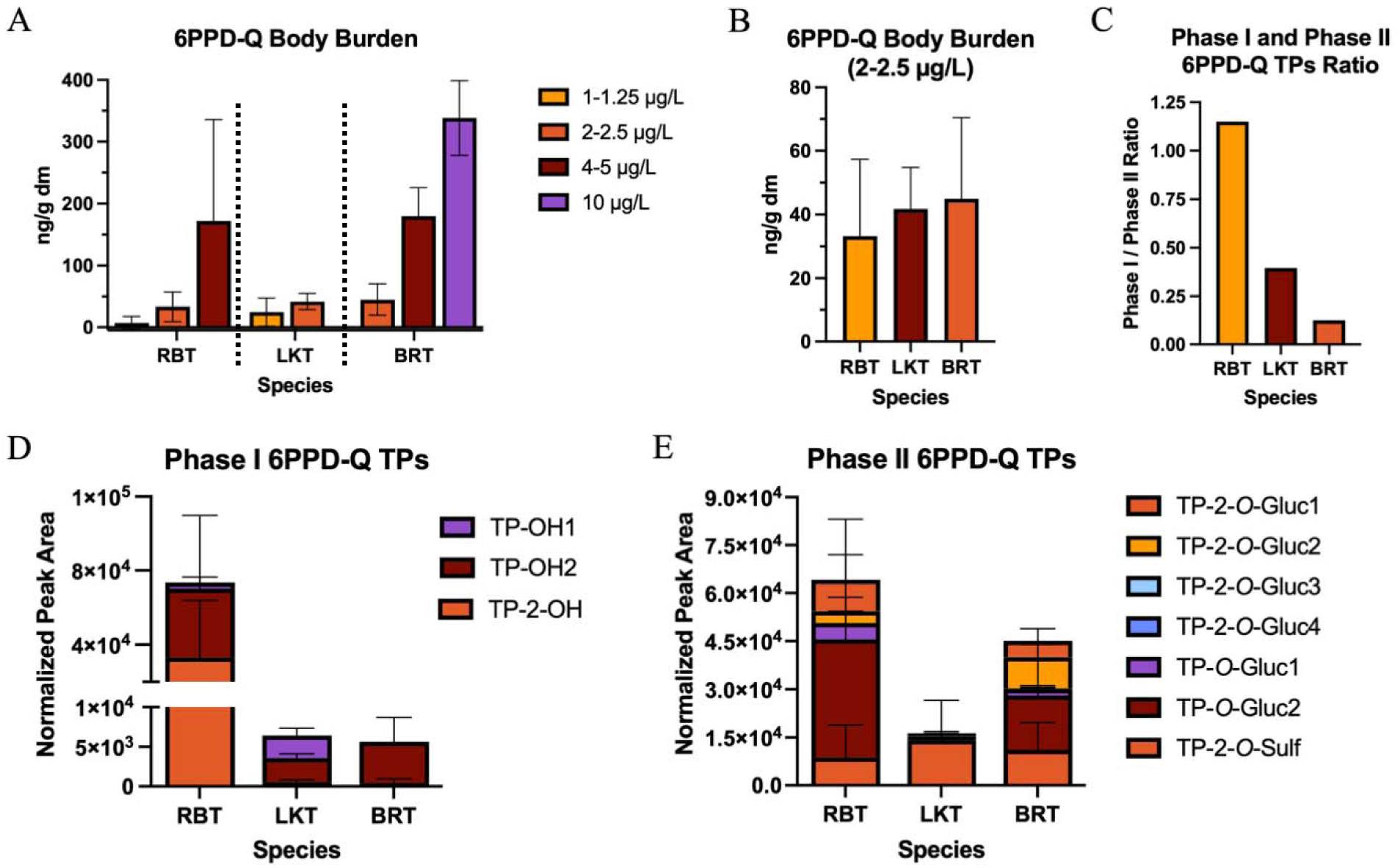
Concentrations and normalized peak areas of 6PPD-Q and phase I and II transformation products (TPs), respectively, in extracts of larval fish tissue. A) Concentrations of 6PPD-Q (ng/g) in extracted fish tissue within nominal exposure bins for lake trout (LKT), rainbow trout (RBT), and brown trout (BRT). B) Concentrations of 6PPD-Q (ng/g) in extracts of fish tissue for 2-2.5 µg/L nominal exposure. C) Ratio of the summed normalized peak areas for phase I and II transformation products (TPs) in 2-2.5 µg/L exposed fishes. D) Phase I 6PPD-Q TPs normalized peak areas in 2-2.5 µg/L exposed fishes. E) Phase II TPs normalized peak areas in 2-2.5 µg/L exposed fishes.

### Detection and Comparison of 6PPD-Q Transformation Products Among Fishes

TPs from biotransformation pathways were detected using suspect screening in 6PPD-Q-exposed larval fish tissue. Three phase I TPs from CYP (Cytochromes P450) biotransformation^13,35^ were detected in fish exposed to 2-2.5 µg 6PPD-Q/L. These TPs included two monohydroxylated TPs and one dihydroxylated TP (Figure 2D). The two monohydroxylated TPs (*m/z* = 315.17) included an alkyl sidechain hydroxylation (TP-OH1, RT = 13.45) and a phenyl ring hydroxylation (TP-OH2, RT = 14.56), both of which have been previously described.^13^ TP-OH1 was verified to be hydroxylated on the carbon 4-position of the alkyl sidechain using a synthesized standard (6-PPD-Q-4-OH, Cayman Chemical),^36^ and TP-OH2 was additionally verified to occur on para position of the phenyl ring using an analytical standard (*p*-hydroxy 6PPD-Q, Cayman Chemical item number 40605) (Supplementary Text S3; Supplementary Figure S8). TP-OH1 was only detected in the more sensitive fish species for the 2-2.5 µg/L exposure bin (Figure 1D), with BRT only having the more abundant TP-OH2 present.^13,37^ The dihydroxylated TP (*m/z* = 331.17) showed identical fragmentation patterns for all species, indicating a phenyl ring hydroxylation and alkyl sidechain hydroxylation, and was found to be greatest in RBT (averaged sum; 3.30×10^4^ arbitrary units, au), followed by LKT (5.74 × 10^2^ au) and BRT (4.31 × 10^2^ au) (Figure 1D). RBT had the greatest amounts of phase I TPs (7.36×10^4^ au; 53.5% peak area of all summed TPs) compared to both LKT (6.44 × 10^3^ au; 28.3%) and BRT (5.64 × 10^3^ au; 11.1%) (Figure 1D). Two additional phase I phenyl ring monohydroxylated TPs were detected in BRT exposed to 10 µg/L. Having a similar fragmentation profile as the phenyl ring monohydroxylation suggests that hydroxylation might have occurred at positions other than the para position (i.e., ortho and meta) at these higher concentrations, leading to different log *K*_ow_ and retention times (RT = 16.48 and 17.83) (Supplementary Figure S1). This provided additional evidence that the preferred phenyl ring monohydroxylation occurred at the para position due to increased availability for CYP biotransformation and decreased steric hindrance.^11^ Phase I to phase II normalized relative abundance ratio values indicated that RBT had the lowest capacity to conjugate hydroxylated TPs (Figure 1C), followed by LKT. BRT had the greatest ability to biotransform hydroxylated TPs further through phase II conjugation, with the phase 1/ phase 2 peak area ratio being lowest for BRT, suggesting overall biotransformation of 6-PPDQ is likely important in *Salmonidae*-specific acute toxicity,^11^ which could have important implications for differences in reported species-specific sensitivities to 6PPD-Q. Affinity for P450 enzymes within *Salmonidae* could play an important role in the enhanced ability of specific species to detoxify 6PPD-Q.^38^ Fish species ability to biotransform chemicals can have important implications in respective sensitivity and can drive changes in endemic biodiversity assemblages.^39^ There is likely a difference between salmonids ability to detoxify 6PPD-Q driving differences in sensitivity, whereas other phylogenetic groups could also have differences in detoxication pathways but additionally changes in target protein structure.^7^

**Figure 2.**
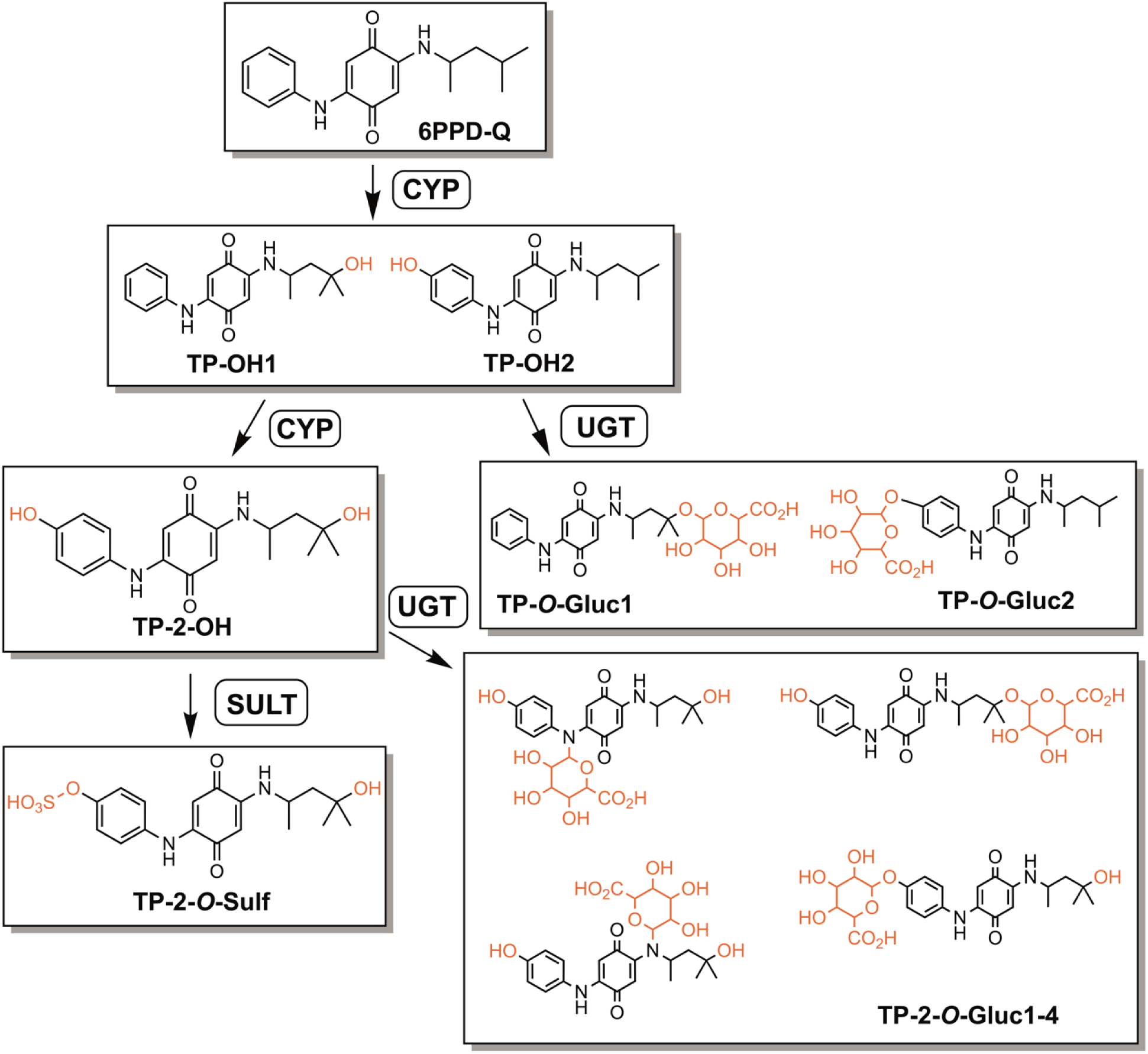
Biotransformation pathways for 6PPD-Q phase I and phase II TPs. 6PPD-Q is biotransformed by CYP to form two different monohydroxylated TPs, TP-OH1 and TP-OH2. Monohydroxylated 6PPD-Q can then undergo glucuronide conjugation *via* UDP-glucuronosyltransferase (UGT) to form two TPs (TP-*O*-Gluc1 and TP-*O*-Gluc2). 6PPD-Q can be further biotransformed by CYP to form a dihydroxylated TP (TP-2-OH). The dihydroxylated TP can be further biotransformed by sulfate (Sulfotransferase; SULT) or glucuronide conjugation (UGT). The probable location of the sulfate conjugation is on the phenyl ring hydroxylation and does not occur on the alkyl sidechain hydroxylation (TP-2-*O*-Sulf; Supplementary Figure S4). Glucuronide conjugation of the dihydroxylated TP can form four different TPs by *N*-glucuronidation or *O*-glucuronidation (TP-2-*O*-Gluc1-4).

A total of seven phase II TPs were detected for 2-2.5 µg/L exposure bin, with presence and relative amounts differing among salmonids (Figure 1E). The seven phase II TPs included two glucuronide-conjugated monohydroxylated TPs (*m/z* = 491.202), four glucuronide-conjugated dihydroxylated TPs (*m/z* = 507.198), and one sulfonate-conjugated dihydroxylated TP (*m/z* = 411.122). In total, five, three, and seven phase II TPs were detected for RBT, LKT, and BRT, respectively (Figure 1D). Greater number of phase II TPs were also detected in the more tolerant zebrafish, a *Cyprinidae*, reflecting a greater capacity to detoxify.^12^ Glucuronide-conjugated TPs dominated in both RBT (5.53 × 10^4^ au; 40.2%) and BRT (3.94 × 10^4^ au; 67.0%), while sulfonate-conjugated TP dominated in LKT (1.24 × 10^4^ au; 62.6%), indicating differences in the biotransformation pathways that predominate among these three salmonids. The most predominant TPs in mammalian models were also monohydroxylated 6PPD-Q and glucuronide-conjugated 6PPD-Q,^40^ similar to both RBT and BRT. RBT had the overall greatest amounts of phase II TPs (6.40 × 10^4^ au; 46.5%), followed by BRT (4.50 × 10^4^ au; 88.9%), with LKT having the lowest relative amounts of phase II TPs (1.63 × 10^4^ au; 71.7%). However, RBT was the only species where phase II TP peak areas were in deficit compared to phase I peak areas (Figure 1C).

The two glucuronide-conjugated monohydroxylated TPs were both *O*-glucuronidated at either the alkyl sidechain hydroxylation (TP-*O*-Gluc1) or phenyl ring hydroxylation (TP-*O*-Gluc2) position (Figure 2, Supplementary Figure S2) with matching fragmentation (MS^2^) to the two monohydroxylated TPs after neutral loss of glucuronide acid (*m/z* = 176.032). The four glucuronide-conjugated dihydroxylated TPs could be either *N*-glucuronidation or *O*-glucuronidation, attributed to four different TPs (Figure 2, Supplementary Figure S3). The single sulfonate-conjugated dihydroxylated TP was likely *O*-sulfonated at the phenyl ring hydroxylation, with an MS^2^ fragment of *m/z* = 311.034 indicating loss of water and an alkyl sidechain (Figure 2, Supplementary Figure S4).

### Metabolome Response of 6PPD-Q-Exposed Fishes

Differences between the greater exposure concentrations and solvent control for RBT and LKT were discerned using PLS-DA (Figure 3A, B), reflecting alterations in the metabolome due to 6PPD-Q exposure. No significant differences were detected for BRT, as all treatments overlapped in 95% confidence intervals (Supplementary Figure S5). RBT and LKT treatments were largely separated by component 1 of the PLS-DA, accounting for 26.5 and 49.1 % of the variability, respectively. Volcano plot for the 2-2.5 µg/L exposure bin, relative to solvent control, revealed a total of 153 (146 increased in the 6PPD-Q treatment group and seven decreased compared to the solvent control) and 170 (nine increased and 161 decreased) dysregulated metabolites for RBT and LKT, respectively (Figure 3C, D), with no dysregulated metabolites for BRT. The shift in dysregulation of metabolites between RBT and LKT could reflect the differences in exposure length between the species (28 d vs. 45 d), with differences in the initial and subsequent compensatory responses. 6PPD-Q toxicity is observed to have differences between acute and sub-chronic toxicity, with a longer duration for LKT potentially driving differences in dysregulated metabolites compared to RBT.^8^ LKT was shown to be slightly more sensitive to 6PPD-Q-exposure, with a 45-d LC_50_ of 0.39 µg/L compared to RBT 28-d LC_50_ of 0.56 µg/L,^8, 23^ which may reflect compensatory mechanisms towards the chemical stressor for RBT and the greater number of increased metabolites.

**Figure 3.**
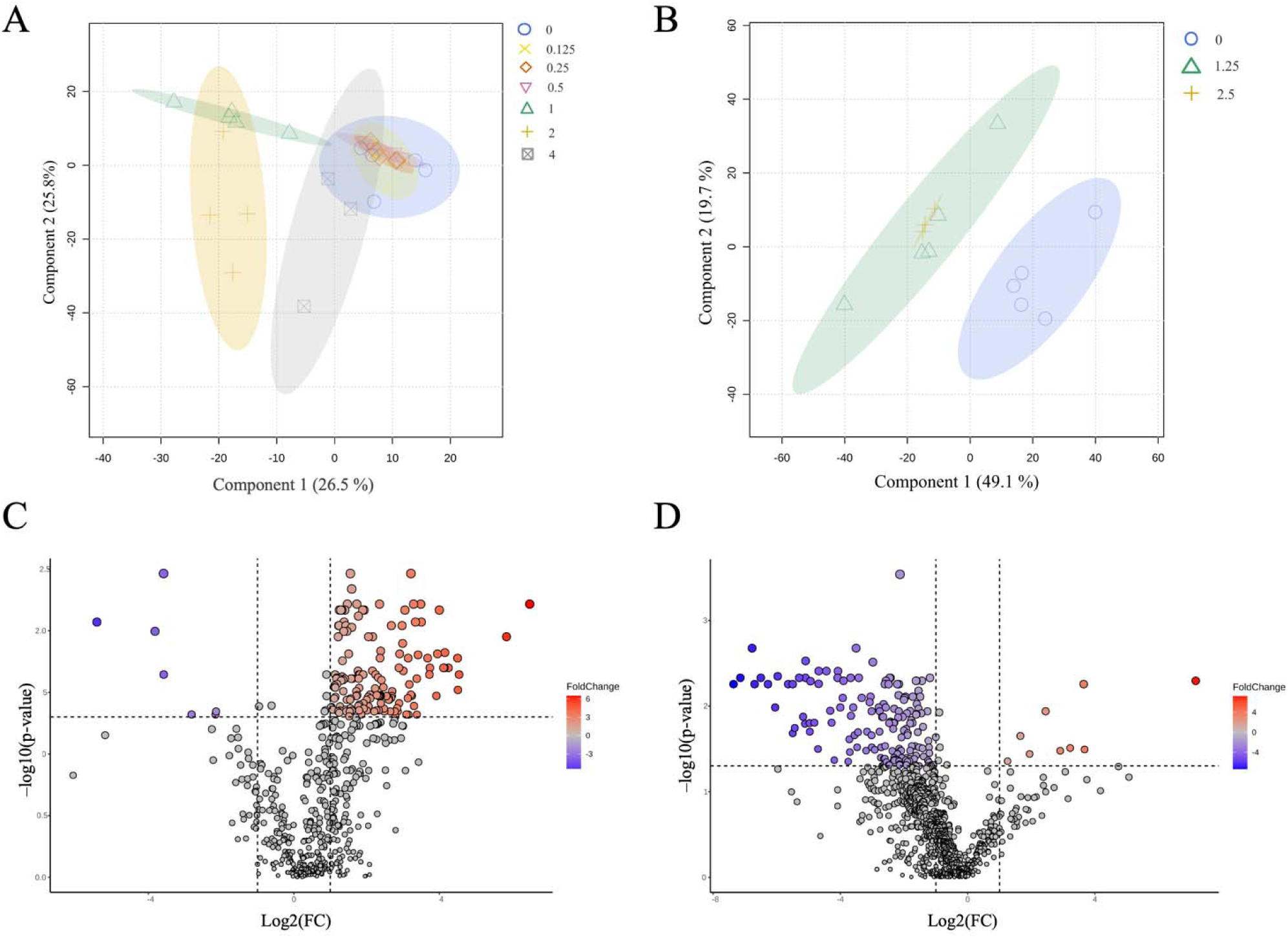
RBT and LKT metabolome responses to 6PPD-Q. PLS-DA of A) RBT metabolome and B) LKT metabolome response to 6PPD-Q. Individual points represent each sample collected. Shapes and colors represent the different treatment groups. Volcano plot of dysregulated metabolites for C) RBT and D) LKT. A false-discovery rate (FDR) *p*-value of 0.05 and fold change (FC) of 2.0 was used for discerning dysregulated metabolites. A negative fold change is displayed in blue, and a positive fold change is displayed in red.

Xanthine, an alkaloid, was the annotated metabolite with the greatest enrichment between solvent control and 6PPD-Q exposure group in RBT (log_2_FC = 6.46; Supplementary Figure S6A; Supplementary File S2), while xanthine was depleted in exposed LKT (log_2_FC = –6.78; Supplementary Figure S6B). Xanthine has a wide array of functions within organisms while also being imperative in the catabolism of nucleotides and nucleic acids, being a precursor of uric acid,^41^ indicating response to stress in RBT and disruption from stress in LKT. Xanthine is also an antagonist of adenosine receptors,^42^ with adenosine having the greatest enrichment in exposed LKT (log_2_FC = 7.17), indicating a balanced relationship between the two metabolites. Adenosine is an endogenous modulator of neuronal excitability,^43^ and the increased abundance of the metabolite could be potentially tied to some of the Urban Runoff Mortality Syndrome (URMS)-related symptoms (e.g., pectoral fin splaying and circling).^44^ Acute toxicity from 6PPD-Q can lead to behavioral abnormalities in exposed salmonids, including loss of equilibrium, erratic swimming patterns, lethargy, and disorientation, with these phenotypes being designated as URMS.^44^ Several additional URMS behaviors from acute 6PPD-Q toxicity are also potentially tied to loss of respiration capability (e.g., surface swimming and mouth gaping).^45^ Several acylcarnitines of mid- to long-chain length were increased (*n* = 6) in RBT from 6PPD-Q exposure, including stearoylcarnitine, valeryl-carnitine, 3-hydroxyhexadecanoylcarnitine, 3-hydroxyhexanoylcarnitine, 3-hydroxyoctadecanoylcarnitine, and butyrylcarnitine, indicating mitochondrial fatty acid oxidation was impacted within this species (perhaps indicative of uncoupling of the mitochondrial electron transport chain).^46,47^ Acylcarnitines are fatty acid metabolites that transport acyl groups from the cytosol into the mitochondrial matrix for *ß*-oxidation and subsequent energy production^48^ and increases in acylcarnitines can be due to mitochondrial dysfunction and alterations in metabolism. LKT had two acylcarnitines decreased in abundance, oleoylcarnitine and L-acetylcarnitine. L-acetylcarnitine, a short-chain acylcarnitine, has a wide range of neuroprotective and neurotrophic effects^48,49^ while also facilitating the transfer of acetyl-Coenzyme A across mitochondrial membranes for energy production with a decrease in L-acetylcarnitine indicating mitochondrial and neuroprotection dysfunction.

Increases in both lysophosphatidylcholines (LPCs; *n* = 6) and phosphatidylcholines (PCs; *n* = 2) were observed in RBT with decreases observed in LKT indicating that there was a disruption in phospholipid biosynthesis, leading to potentially altered membrane permeability.^50^ Higher concentrations of LPCs can also disrupt mitochondrial integrity and modulate inflammatory chemokine expression.^51,52^ Changing levels of PCs can lead to neurotoxicity from altered levels of acetylcholine leading to additional neurological effects.^53,54^ Greater amounts of dysregulated LPCs derived from the cleaving of PC indicate activation of signaling pathways that are involved in oxidative stress and inflammatory responses.^52^ Disruption in blood-brain barrier (BBB) permeability was an observed transcriptome response in developing coho salmon that was indicated to alter central nervous system function.^55^ LPCs are important for the transport of polyunsaturated fatty acids (PUFAs) across BBB via major-facilitator superfamily transporter MFSD2A.^56^ Several long-chain polyunsaturated fatty acids (LC-PUFAs) were seen to increase in sub-chronically exposed RBT including arachidonic acid (AA; omega-6 fatty acid), 8,9-epoxyeicosatrienoic acid (a metabolite of AA formed through P450 epoxygenases), Dihomo-gamma-linolenic acid (omega-6 fatty acid), docosahexaenoic acid (DHA; omega-3 fatty acid), eicosapentaenoic acid ethyl ester (omega-3 fatty acid that is an esterified form of eicosapentaenoic acid (EPA)), and docosapentaenoic acid (DPA; omega-3 fatty acid that is an intermediate between EPA and DHA). Increases in these LC-PUFAs could indicate alterations in the AA pathway, with importance to inflammatory processes and disease,^57^ while alterations in important omega-3 PUFAs could modify key involvement in anti-inflammatory processes and cell membrane permeability.^58^ Several LC-PUFAs were observed to decrease in exposed LKT including AA, DPA, EPA, linoleic acid, and dihomo-alpha-linolenic acid indicating deficiencies in essential fatty acids that could lead to altered metabolic and membrane function, chronic inflammation, neurological dysregulation, and oxidative stress.^59^ Additional metabolites with the greatest increased FC in exposed RBT included 11-hydroxy DHA (log_2_FC = 4.51; Supplementary Figure S6C), a marker of inflammation and the result of oxidation,^60^ palmitoleic acid (log_2_FC = 4.10; Supplementary Figure S6D), 14S-hydroxy-4Z,7Z,10Z,12E,16Z,19Z-docosahexaenoic acid (log_2_FC = 4.08), an additional marker of oxidative stress resulting from oxidation of DHA,^61^ and inosine (log_2_FC = 3.51), a nucleoside and versatile bioactive metabolite with increases in abundance linked to regulating inflammation, immunomodulatory effects, and neuroprotection.^62,63^

No increased or decreased metabolites overlapped between RBT and LKT, indicating that these two species differed in their physiological response to 6PPD-Q at the time of sampling. Results indicate that RBT were trying to manage stressor impacts further by compensation with a shorter exposure time, whereas LKT physiological processes were ultimately deteriorating from an extended exposure duration, reflecting the differences in the biotransformation capability between the two species. More effort will be needed to profile endogenous metabolite alterations, including temporal responses, using developed target LC-HRMS methods for specific metabolite classes. Additional research is also needed to link changes in metabolome profiles directly with URMS behaviors and neurotoxicity.

Using dose-response analysis, an mPOD was computed for the RBT metabolome (Supplementary Figure S7), while LKT and BRT metabolome data were not conducive to mPOD analysis due to the limited number of doses or a low number of dysregulated metabolites, respectively.^64,65^ A total of 213 metabolites were fitted with models, with 203 metabolites having BMDs for mPOD computation. The RBT mPOD had a 0.44 µg/L feature-level BMD for the 20^th^ feature and 10^th^ percentile, which closely aligns with the 28-d LC_50_ of larval RBT and is protective of the adult RBT 24-h LC_50_ of 1.96 µg/L or 72-g LC_50_ = 1.00 µg/L.^7^ Dose-response analysis of metabolomics datasets is an emerging area of research in aquatic toxicology that could be widely indicative of aquatic health through predictive means and used for chemical risk assessment,^66^ as portrayed here using 6PPD-Q as a study chemical. Utilizing generated non-targeted metabolomics datasets can provide an efficient method for calculating mPODs from organismal response to chemicals.

The xenometabolome of three different salmonids exposed to 6PPD-Q was profiled, revealing that three phase I and seven phase II TPs dominated at lower exposure to 6PPD-Q. Different biotransformation pathways can dominate among salmonids, with RBT and BRT having greater levels of glucuronide-conjugated TPs than the sulfate-conjugated TP found to dominate in LKT. This can have important implications regarding species-specific biotransformation and subsequent response to contaminants, including 6PPD-Q. More research is needed to understand the main mechanisms of toxicological responses to 6PPD-Q and its phase I TPs, including TP-OH1, which was detected in both RBT and LKT but not BRT. Synthesized analytical standards were used to verify the structure of both TP-OH1 and TP-OH2 and will be useful for future biomonitoring and exposure assessment. A time-course metabolome response to 6PPD-Q is also needed to understand how larval fish initially respond to the contaminant and subsequent compensatory responses to have a greater mechanistic understanding of 6PPD-Q toxicity. These results indicate that a difference in biotransformation capability and pathways is observed among salmonid fish species, and subsequent metabolome response to 6PPD-Q exposure can help reveal the mode of toxic action.

## Acknowledgment

This project was supported partially by a financial contribution from Fisheries and Oceans Canada. Additional funding was provided to M.H., S.W., N.H., and M.B. through the Discovery Grants program of the Natural Sciences and Engineering Research Council of Canada (NSERC). M.H. was also supported by the Canada Research Chairs program. Instrumentation used for chemical analyses within this research was obtained with funding from Western Economic Diversification Canada (WED) and the Canadian Foundation for Innovation (CFI). P.J.A. was supported through a Toxicology Centre Devolved Scholarship. M.B. is currently a faculty member of the Global Water Futures (GWF) program, which received funds from the Canada First Research Excellence Funds (CFREF). The graphical abstract was created with Bioicons.com. Additional support came from the staff at Allison Creek Brood Trout Station, who provided the Brown trout embryos, as well as the technical staff at the ARF at the University of Lethbridge (Dr. S. Mamun and H. Shepherd).

