## Supplemental Information for "Xenometabolome of Early-Life Stage Salmonids Exposed to 6PPD-Quinone"

**CONTENT**

**Supplementary Text S1.** Targeted analysis for exposure verification using LC-HRMS.

**Supplementary Text S2.** Non-targeted analysis of larval fish xenometabolome using LC-HRMS.

**Supplementary Text S3.** Confirmation of monohydroxylated TPs structure.

**Supplementary Text S4.** Synthesis of 6-PPD-Q-OH standards.

**Supplementary Text S5.** Filtering of metabolome data for statistical analysis in MetaboAnalyst.

**Supplementary Figure S1.** Phenyl ring monohydroxylation TPs in BRT probable structure and MS^2^.

**Supplementary Figure S2.** 6PPD-Q glucuronide-conjugated monohydroxylated TPs structure and MS^2^.

**Supplementary Figure S3.** 6PPD-Q glucuronide-conjugated dihydroxylated TPs structure and MS^2^.

**Supplementary Figure S4.** 6PPD-Q sulfonate-conjugated dihydroxylated TP structure and MS^2^.

**Supplementary Figure S5.** PLS-DA of BRT metabolome response to 6PPD-Q exposure.

**Supplementary Figure S6.** Violin plots of select dysregulated metabolites in 6PPD-Q exposed fish.

**Supplementary Figure S7.** Metabolomics points-of-departure (mPOD) of RBT exposed to 6PPD-Q.

**Supplementary Figure S8.** Retention time and MS^2^ fragmentation of 6-PPD-Q-4-OH, 6PPD-Q-5-OH, 6PPD-Q-1-OH, *p*-hydroxy 6PPD-Q (Cayman Chemical), TP-OH1, and TP-OH2.

**Supplementary Table S1.** Experimental treatment groups, nominal concentrations of 6PPD-Q exposure, and larval fish tissue dry weight.

**Supplementary Table S2.** Water chemistry.

**Supplementary Table S3.** Positive mode targeted gradient elution method.

**Supplementary Table S4.** Positive mode suspect screening gradient elution method.

**Supplementary Table S5.** Precursor and product ions ([M+H]+) and retention time details for 6PPD-Q, 6PPD-Q-d5, Phase I, and Phase II transformation products.

**Supplementary Text S1.**

Targeted analysis of 6PPD-Q for aqueous exposure verification was performed using a Vanquish UHPLC paired with the Q-Exactive^TM^ high-field (HF) hybrid Quadrupole-Orbitrap mass spectrometer (MS) (Thermo-Fisher), LC separation was achieved using a Kinetex® 1.7 µm XB-C18 column (100 x 2.1 mm). The aqueous and organic mobile phases consisted of 5 % methanol in water with 0.1 % formic acid and 100 % methanol with 0.1 % formic acid, respectively. The flow gradient of the mobile phases is found in Supplementary Table S3. Sample injection volumes of 5.0 µL were ionized using heated electrospray ionization (HESI) in positive mode with a scan range of 100 – 1000 m/z. Ion source parameters were as follows: sheath gas flow = 35; aux gas flow = 10; sweep gas flow = 1; spray voltage = 4.00 kV; capillary temperature = 350 °C; S-lens RF level = 60; and aux gas heater temperature = 300 °C. A paired full scan and parallel reaction monitoring (PRM) MS method was used, with max injection times = 100 ms, resolutions of 120,000/30,000, and AGC targets of 1.0x10^6^/2.0x10^5^, respectively. Transition ions of 6PPD-Q (*m/z* = 215.082) and 6PPD-Q d5 (*m/z* = 220.113) were used for quantification. The instrumental analysis of 6PPD-Q included a 7-point calibration curve, ranging from 0.05 – 100.0 µg/L with an r^2^ = 0.9995 using a weighting factor of 1/X for quantitation. Instrument blanks and QA/QC standards were run once after every nine samples to monitor for carryover and instrument drift. Data was quantified using TraceFinder (version 4.1). The method detection limit (MDL = σ * t_(n-1, 1-α=0.99)_),^1^ and limits of detection (LOD = 3.9 * (σ_pseudo-blank_/slope)) and quantitation (LOQ = 3.3 * LOD)^2^ were 0.027, 0.0080, and 0.026 µg/L, respectively.

**Supplementary Text S2.**

LC separation of analytes was achieved with a Kinetex 1.7-μm XB-C18 LC column (100 × 2.1 mm) (Phenomenex, CA) using Vanquish UHPLC with gradient elution (Supplementary Table S4) and flow rate = 0.2 mL/min, column temperature = 40 °C, solvent A = 95% H2O: 5% MeOH + 0.1% formic acid, and solvent B = 100% MeOH + 0.1% formic acid.

**Supplementary Text S3.**

Targeted analysis was performed using a Vanquish UHPLC paired with the Q-Exactive^TM^ high-field (HF) hybrid Quadrupole-Orbitrap mass spectrometer (MS) (Thermo-Fisher), LC separation was achieved using a Kinetex® 1.7 µm XB-C18 column (100 x 2.1 mm). The aqueous and organic mobile phases consisted of 5 % methanol in water with 0.1 % formic acid and 100 % methanol with 0.1 % formic acid, respectively. The flow gradient of the mobile phase was the same as the non-targeted run (Supplementary Table S4). Sample injection volumes of 10.0 µL were ionized using heated electrospray ionization (HESI) in positive mode with a scan range of 100 – 1000 m/z. Ion source parameters were as follows: sheath gas flow = 35; aux gas flow = 10; sweep gas flow = 1; spray voltage = 4.00 kV; capillary temperature = 350 °C; S-lens RF level = 60; and aux gas heater temperature = 300 °C. A paired full scan and parallel reaction monitoring (PRM) MS method was used, with max injection times = 100 ms, resolutions of 120,000/30,000, and AGC targets of 1.0x10^6^/2.0x10^5^, respectively, with nominal collision energy (NCE) set to 30. 6-PPD-Q-4-OH, 6-PPD-Q-5-OH, 6-PPD-Q-1-OH (item number 41591), and *p*-hydroxy 6-PPD-Q (item number 40605) were run at 10 µg/L dissolved in 50:50 Methanol/Water. Information regarding the synthesis of these standards can be found in Supplementary Text S4. Along with these standards, immortalized RTL-W1 cells incubated with varying concentrations of 6PPD-Q (10 and 160 µg/L) for 24 h media were analyzed to verify the phase I transformation products following methods outlined previously.^3^

6-PPD-Q-4-OH, 6-PPD-Q-5-OH, 6-PPD-Q-1-OH, and *p*-hydroxy 6-PPD-Q had retention times of 13.23 min, 13.26 min, 14.23 min, and 14.32 min (Supplementary Figure S8). 6-PPD-Q-4-OH had MS^2^ fragments with greatest relative abundance of *m/z* 187.087, 200.071, 215.082, 241.097, 257.129, and 297.160. 6-PPD-Q-5-OH had MS^2^ fragments with greatest relative abundance of *m/z* 99.081, 187.087, 200.071, 215.082, 241.097, and 297.160. 6-PPD-Q-1-OH had MS^2^ fragments with greatest relative abundance of *m/z* 187.087, 200.071, 216.089, and 241.097. *p*-hydroxy 6-PPD-Q had MS^2^ fragments with greatest relative abundance of *m/z* 203.082, 231.077, and 272.116.

Confirmation of 6-PPD-Q-4-OH with TP-OH1 with both having a retention time of 13.23 min was observed in the cell media samples with the appropriate fragmentation (*m/z* = 187.087, 200.071, 215.082, 241.097, 257.129 and 297.160) and relative abundance of the fragments (Supplementary Figure S8). Confirmation of *p*-hydroxy 6-PPD-Q with TP-OH2 with a shared retention time of 14.33 min was observed in the cell media samples with the appropriate fragmentation (*m/z* = 203.082, 231.076, and 272.116) and relative abundance of the fragments (Supplementary Figure S8). Both 6-PPD-Q-5-OH and 6-PPD-Q-1-OH were determined not to be the structure of the alkyl sidechain monohydroxylation transformation product (TP-OH1). 6-PPD-Q-5-OH had a similar retention time compared to TP-OH1 (13.23 min) and had confirming MS^2^ fragments of *m/z* 187.087, 200.071, 215.082, 241.097, and 297.160 (Supplementary Figure S8). However, there were some discrepancies between the two analytes, with 6-PPD-Q-5-OH missing the MS^2^ fragment 257.129 found for TP-OH1 along with the slight shift in retention time (13.26 min). 6-PPD-Q-1-OH had distinguishing fragments from TP-OH1, including MS^2^ fragments *m/z* 215.082 and 257.128, along with the large shift in retention time (14.23 min vs 13.23).

**Supplementary Text S4.**

***Synthesis of 6-PPD-Q-5-OH (1)***

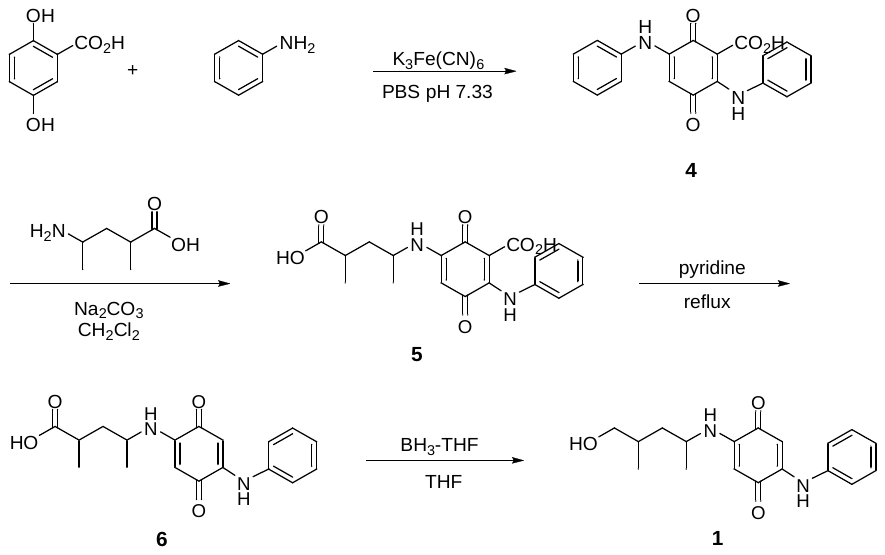

**3,6-dioxo-2,5-bis(phenylamino)cyclohexa-1,4-diene-1-carboxylic acid (4)** To a stirred solution of 2,5-dihydroxybenzoic acid (2.0 g, 12.98 mmol) in 120 mL phosphate buffer (pH 7.33) was added potassium ferricyanide (8.55 g, 25.96 mmol) followed by aniline (3.63 g, 38.94 mmol). The reaction was stirred at room temperature for 18 h. The reaction mixture was extracted with CH_2_Cl_2_ (1L), then washed with 1N HCl (2 x 200 mL) and brine (2 x 200 mL). The organic solution was dried over Na_2_SO_4_, filtered, and evaporated *in vacuo*. The crude product **4** was obtained as a light brown solid and used without further purification.

**5-((4-carboxypentan-2-yl)amino)-3,6-dioxo-2-(phenylamino)cyclohexa-1,4-diene-1-carboxylic acid (5)** To a stirred solution of crude compound **4** (160 mg, 0.479 mmol) in 10 mL CH_2_Cl_2_ was added Na_2_CO_3_ (51 mg, 0.479 mmol). 4-amino-2-methylpentanoic acid (80 mg, 0.479 mmol) was pre-treated with 0.1M NaOH in MeOH (4.79 mL, 1.0 eq) in a separate flask, then evaporated to dryness, dissolved in 1 mL CH_2_Cl_2_, and added dropwise to the reaction mixture. The reaction was stirred at room temperature for 72 h. The reaction mixture was diluted with CH_2_Cl_2_ (50 mL) and washed with 1N HCl (2 x 10 mL) and brine (2 x 10 mL). The organic solution was dried over sodium sulfate, filtered, and evaporated *in vacuo*. The crude product was purified by flash column chromatography (SiO_2_; CH_2_Cl_2_/MeOH/AcOH 10:0:0.1 to 9.8:0.2:0.1, v/v) to afford **5,** a mixture of diastereomers, (133 mg, 75%) as a red solid. MS (dual ESI/APCI) *m/z* calculated for C_19_H_20_N_2_O_6_ [M-H]^-^ 371.13, found 371.1.

**4-((3,6-dioxo-4-(phenylamino)cyclohexa-1,4-dien-1-yl)amino)-2-methylpentanoic acid (6)**

A solution of **5** (133 mg, 0.357 mmol) in 15 mL of pyridine was refluxed at 120°C for 18 h. The reaction mixture was cooled to room temperature and concentrated *in vacuo*. The residue was dissolved in CH_2_Cl_2_ (100 mL) then washed with 0.1N HCl (3 x 25 mL) and brine (1 x 25mL). The organic solution was dried with Na_2_SO_4_, filtered, and evaporated *in vacuo.* The crude product was purified by flash column chromatography (SiO_2_; heptane/EtOAc 10:0 to 7:3, v/v) to afford **6,** a mixture of diastereomers, (27 mg, 23%) as a red solid. ^1^H NMR (400 MHz, Chloroform-d) δ ppm 1.18-1.34 (m, 6H), 1.52-1.75 (m, 1H), 1.96-2.19 (m, 1H), 2.47-2.65 (m, 1H), 3.52-3.74 (m, 1H), 5.41-5.52 (m, 1H), 5.94 (m, 1H), 6.40-6.54 (m, 1H), 7.16-7.27 (m, 3H), 7.35-7.43 (m, 2H), 8.12-8.32 (m, 1H). MS (dual ESI/APCI) *m/z* calculated for C_19_H_20_N_2_O_6_ [M-H]^-^ 327.14, found 327.1.

**2-((5-hydroxy-4-methylpentan-2-yl)amino)-5-(phenylamino)cyclohexa-2,5-diene-1,4-dione (1)** To a stirred solution of compound **6** (13 mg, 0.0396 mmol) in 1 mL THF was added 1.0 M Borane-THF (1:1) (59 uL, 0.0594 mmol) dropwise at 0°C. The mixture was stirred at room temperature for 18 h. DI H_2_O was added, then extracted with EtOAc (3 x 25). The combined organic layers were washed with 1M HCl (2 x 15 mL), DI H_2_O (2 x 15 mL), and brine (2 x 15 mL). The organic solution was dried over Na_2_SO_4,_ filtered, and evaporated *in vacuo.* The crude product was purified by flash column chromatography (SiO_2_; CH_2_Cl_2_/MeOH/AcOH 10:0:0.1 to 9.8:0.2:0.1, v/v) to afford **1**, a mixture of diastereomers, (8.6 mg, 69%) as a red solid. ^1^H NMR (400 MHz, Chloroform-d) δ ppm 0.89-1.00 (m, 3H), 1.16-1.32 (m, 3H), 1.33-1.44 (m, 1H), 1.51 (dt, *J*=13.98, 7.22 Hz, 1H), 1.60-1.86 (m, 2H), 3.43-3.54 (m, 2H), 3.58-3.65 (m, 1H), 5.42-5.45 (m, 1H), 5.96 (s, 1H), 6.43-6.56 (m, 1H), 7.18-7.28 (m, 3H), 7.39 (t, *J*=7.56 Hz, 2H), 8.20 (br s, 1H). MS (dual ESI/APCI) *m/z* calculated for C_18_H_22_N_2_O_3_ [M-H]^-^ 313.16, found 313.1.

***Synthesis of 6-PPD-Q-1-OH (2)***

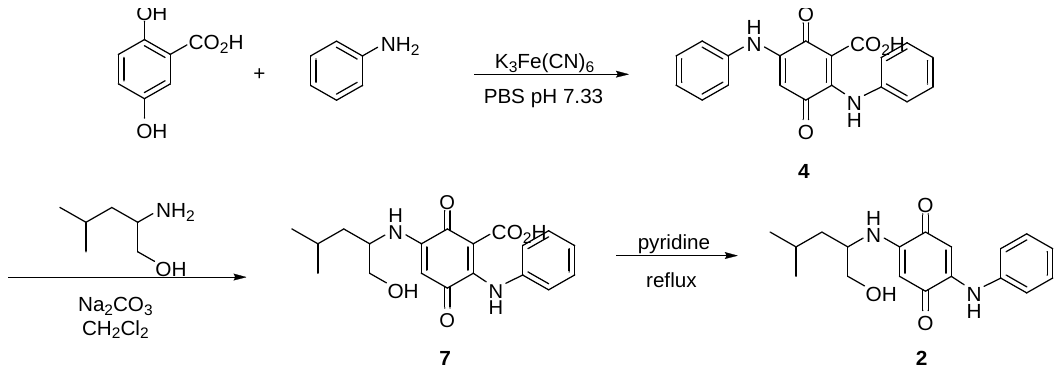

**3,6-dioxo-2,5-bis(phenylamino)cyclohexa-1,4-diene-1-carboxylic acid (4)** To a stirred solution of 2,5-dihydroxybenzoic acid (2.0 g, 12.98 mmol) in 120 mL phosphate buffer (pH 7.33) was added potassium ferricyanide (8.55 g, 25.96 mmol) followed by aniline (3.63 g, 38.94 mmol). The reaction was stirred at room temperature for 18 h. The reaction mixture was extracted with CH_2_Cl_2_ (1 L), then washed with 1N HCl (2 x 200 mL) and brine (2 x 200 mL). The organic solution was dried over Na_2_SO_4_, filtered, and evaporated *in vacuo*. The crude product **4** was obtained as a light brown solid and used without further purification.

**5-((1-hydroxy-4-methylpentan-2-yl)amino)-3,6-dioxo-2-(phenylamino)cyclohexa-1,4-diene-1-carboxylic acid (7)** To a stirred solution of crude compound 4 (175 mg, 0.523 mmol) in 10 mL CH_2_Cl_2_ was added Na_2_CO_3_ (55 mg, 0.523 mmol). 2-amino-4-methyl-pentan-1-ol (61 mg, 0.523 mmol) was dissolved in 1 mL dichloromethane and added dropwise to the reaction mixture. The reaction was stirred at room temperature for 18 h. The reaction mixture was diluted with CH_2_Cl_2_ (50 mL) and washed with 1N HCl (2 x 10 mL) and brine (2 x 10 mL). The organic solution was dried over Na_2_SO_4,_ filtered, and evaporated *in vacuo*. The crude product was purified by flash column chromatography (SiO_2_; CH_2_Cl_2_/EtOAc/AcOH 10:0:0.1 to 9.5:5:0.1, v/v) to afford **7**, a mixture of enantiomers, (174 mg, 93%) as a red oil. ^1^H NMR (400 MHz, Chloroform-d): δ ppm 0.87-0.91 (d, 3H), 0.94-0.98 (d, 3H), 1.43-1.83 (m, 5H), 3.60-3.69 (dd, 1H), 3.83-3.90 (dd, 1H), 5.24-5.38 (m, 1H), 6.04 (s, 1H), 7.21-7.31 (m, 3H), 7.40-7.47 (m, 2H), 8.17 (br s, 1H), 12.36 (br s, 1H). MS (dual ESI/APCI) *m/z* calculated for C_19_H_22_N_2_O_5_ [M-H]^-^ 357.15, found 357.1.

**2-((1-hydroxy-4-methylpentan-2-yl)amino)-5-(phenylamino)cyclohexa-2,5-diene-1,4-dione (2)** A solution of **7** (174 mg, 0.486 mmol) in 20 mL of pyridine was refluxed at 120°C for 5 h. The reaction mixture was cooled to room temperature and concentrated *in vacuo*. The residue was dissolved in CH_2_Cl_2_ (100 mL) and washed with 0.1N HCl (3 x 25 mL) and brine (1 x 25 mL). The organic solution was dried over Na_2_SO_4_, filtered, and evaporated *in vacuo.* The crude product was purified by flash column chromatography (SiO_2_; heptane/EtOAc 10:0 to 7:3, v/v) to afford **2**, a mixture of enantiomers, (62 mg, 41%) as a red solid. ^1^H NMR (400 MHz, Chloroform‑d) δ ppm 0.91-0.92 (d, 3H), 0.94-0.96 (d, 3H), 1.49-1.54 (m, 2H), 1.61-1.71 (m, 1H), 3.53-3.61 (m, 1H), 3.62-3.68 (dd, 1H), 3.72 -3.78 (dd, 1H), 5.50 (s, 1H), 5.97 (s, 1H), 6.50 (br d, J=8.71 Hz, 1H), 7.19-7.23 (m, 3H), 7.35-7.43 (m, 2H), 8.17 (br s, 1H). MS (dual ESI/APCI) *m/z* calculated for C_18_H_22_N_2_O_3_ [M-H]^-^ 313.16, found 313.1.

***Synthesis of p-hydroxy-6-PPD-Q (3)***

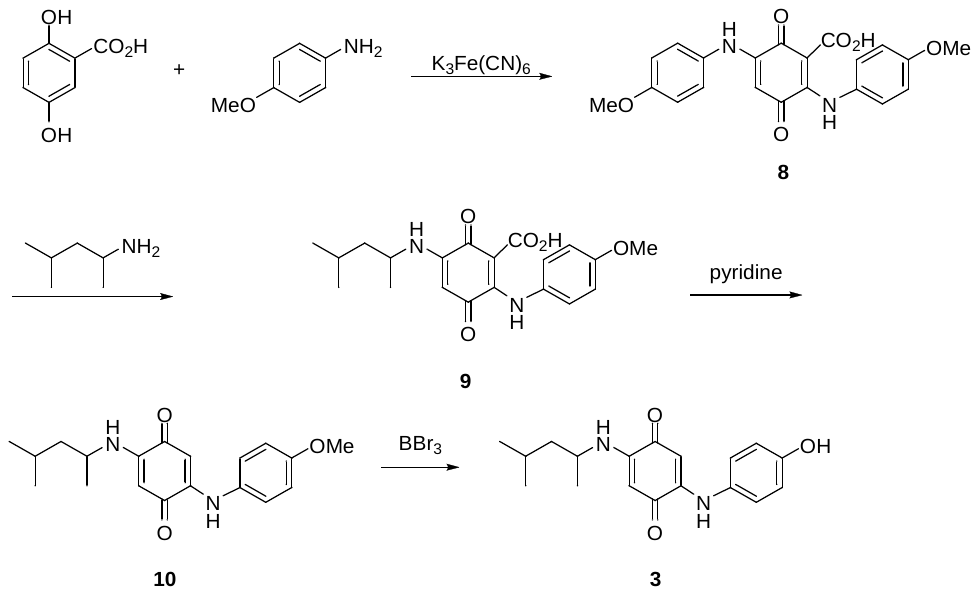

**2,5-bis((4-methoxyphenyl)amino)-3,6-dioxocyclohexa-1,4-diene-1-carboxylic acid (8)** To a stirred solution of 2,5-dihydroxybenzoic acid (292 mg, 1.89 mmol) in 20 mL of phosphate buffer (pH 7.33) was added potassium ferricyanide (1.25 g, 3.79 mmol) followed by p-anisidine (700 mg, 5.68 mmol). The reaction was stirred at room temperature for 18 h. The reaction mixture was extracted with CH_2_Cl_2_ (1 x 150 mL, 2 x 20 mL) then washed with 1N HCl (2 x 20 mL) and brine (1 x 20 mL). The organic solution was dried over Na_2_SO_4_, filtered and evaporated *in vacuo*. The crude product **8** was obtained as a light brown solid and used without further purification.

**2-((4-methoxyphenyl)amino)-5-((4-methylpentan-2-yl)amino)-3,6-dioxocyclohexa-1,4-diene-1-carboxylic acid (9)** To a stirred solution of crude compound **8** (747 mg, 1.89 mmol) in 10 mL CH_2_Cl_2_ was added Na_2_CO_3_ (201 mg, 1.89 mmol). 1,3-Dimethylbutylamine (192 mg, 1.89 mmol) was dissolved in 500 mL CH_2_Cl_2_ and added dropwise to the reaction mixture. The reaction was stirred at room temperature for 18 h. The reaction mixture was diluted with CH_2_Cl_2_ (75 mL) and washed with 1N HCl (2 x 10 mL) then brine (1 x 10 mL). The organic solution was dried over Na_2_SO_4_, filtered and evaporated *in vacuo*. The residue was purified by flash column chromatography (SiO_2_, CH_2_Cl_2_ with 0.5% AcOH) to afford **9**, a mixture of enantiomers, (185 mg, 26%) as a red solid. ^1^H NMR (400MHz, Chloroform-d): d ppm 0.88-0.90 (d, 3H), 0.95-0.96 (d, 3H), 1.33-1.35 (d, 3H), 1.41-1.47 (m, 1H), 1.58-1.72 (m, 3H), 3.85 (s, 3H), 5.17-5.24 (m, 1H), 6.94-6.97 (m, 2H), 7.17-7.20 (m, 3H), 8.03 (s, 1H), 12.33 (s, 1H). MS (dual ESI/APCI) *m/z* calculated for C_20_H_24_N_2_O_5_ [M+H]^+^ 373.17, found 373.2.

**2-((4-methoxyphenyl)amino)-5-((4-methylpentan-2-yl)amino)cyclohexa-2,5-diene-1,4-dione (10)** A solution of **9** in 20 mL pyridine was refluxed at 120°C for 18 h. The reaction mixture was cooled to room temperature and concentrated *in vacuo*. The residue was dissolved in CH_2_Cl_2_ (100 mL) and washed with 0.1N HCl (3 x 10 mL) then brine (1 x 10 mL). The organic solution was dried over Na_2_SO_4_, filtered and evaporated *in vacuo* to afford **10**, a mixture of enantiomers, (134 mg, 97%) as a purple solid. ^1^H NMR (400MHz, Chloroform-d): d ppm 0.89-0.93 (d, 3H), 0.93-0.96 (d, 3H), 1.20-1.22 (d, 3H), 1.34-1.40 (m, 1H), 1.48-1.55 (m, 1H), 1.63-1.68 (m, 1H), 3.51-3.56 (m, 1H), 3.81 (s, 3H), 5.40 (s, 1H), 5.79 (s, 1H), 6.40-6.42 (br d, 1H), 6.90-6.92 (d, 2H), 7.15-7.17 (d, 2H), 8.11 (br s, 1H). MS (dual ESI/APCI) *m/z* calculated for C_19_H_24_N_2_O_3_ [M+H]^+^ 329.18, found 329.2.

***p*-hydroxy-6-PPD-Q (3)** To a stirred solution of **10** in 2 mL CH_2_Cl_2_ at 0°C was added dropwise boron tribromide (3.65 mL, 3.65 mmol). The reaction was stirred at room temperature for 18 h. After cooling to 0°C, 10% aqueous NH_4_OH was added dropwise until pH 8. The reaction mixture was diluted with CH_2_Cl_2_ (200 mL) and washed with brine (1 x 15 mL). The organic solution was dried over Na_2_SO_4_, filtered and evaporated *in vacuo*. The residue was purified by column chromatography (SiO_2_, heptane/EtOAc 10:0 to 7:3, v/v) to afford **3**, a mixture of enantiomers, (62 mg, 54%) as a brown solid. ^1^H NMR (400MHz, Chloroform-d): d ppm 0.87-0.90 (d, 3H), 0.91-0.93 (d, 3H), 1.21-1.23 (d, 3H), 1.33-1.40 (m, 1H), 1.49-1.56 (m, 1H), 1.61-1.68 (m, 1H), 3.51-3.58 (m, 1H), 5.41 (s, 1H), 5.77 (s, 1H), 6.42-6.45 (br d, 1H), 6.84-6.86 (d, 2H), 7.09-7.11 (d, 2H), 8.13 (br s, 1H). MS (dual ESI/APCI) *m/z* calculated for C_18_H_22_N_2_O_3_ [M+H]^+^ 315.16, found 315.2.

***Synthesis of 6-PPD-Q-4-OH (12)***

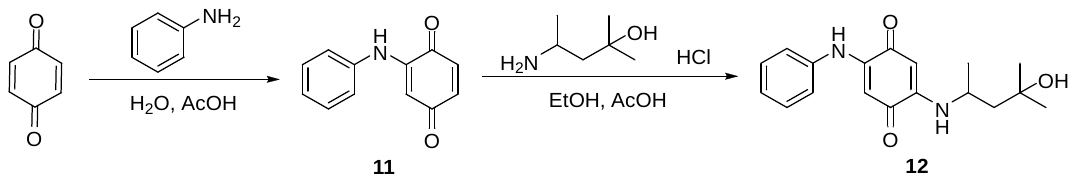

**2-(phenylamino)cyclohexa-2,5-diene-1,4-dione (11)** 1,4-benzoquinone (1.00g, 9.25 mmol) was dissolved in hot water (125 mL), and the solution was cooled to room temperature. Aniline (439 mg, 4.71 mmol) was dissolved in acetic acid (0.25 mL) and water (2.75 mL) and added to the 1,4-benzoquinone solution. The reaction was stirred at room temperature for 18 h. Saturated sodium bicarbonate (25 mL) was added. The resulting mixture was extracted with CH_2_Cl_2_ (3 x 50 mL), then washed with DI H_2_O (3 x 50 mL) and brine (1 x 50 mL). The organic solution was dried over Na_2_SO_4_, filtered, and evaporated *in vacuo*. The solid was purified by flash column chromatography (SiO_2_; heptane/EtOAc 10:0 to 40:60, v/v) to afford **11** (591 mg, 32%) as a brown solid. ^1^H NMR (400 MHz, Chloroform-d): d ppm 6.18 (d, 1H), 6.69-6.72 (m, 2H), 7.17-7.23 (m, 3H), 7.39 (t, *J*=7.61 Hz, 2H)

**2-((4-hydroxy-4-methylpentan-2-yl)amino)-5-(phenylamino)cyclohexa-2,5-diene-1,4-dione (12)** A stirred solution of crude compound 11 (100 mg, 0.502 mmol) in 1.5 mL ethanol was cooled to 0°C. 4-amino-2-methylpentan-2-ol hydrochloride (50 mg, 0.325 mmol) was pre-treated with 0.1M NaOH in MeOH (3.25 mL, 1.0 eq) in a separate flask to desalt the amine. This was evaporated to dryness, dissolved in ethanol (1mL) and acetic acid (0.1 mL), and added dropwise to the reaction mixture at 0°C. The reaction was stirred at room temperature for 18 h. Saturated sodium bicarbonate was added (10 mL), and the aqueous was extracted with EtOAc (3 x 25 mL). The combined organic layers were washed with DI H_2_O (2 x 15 mL) and brine (2 x 15 mL). The organic solution was dried over Na_2_SO_4,_ filtered, and evaporated *in vacuo.* The crude product was purified by flash column chromatography (SiO_2_; heptane/EtOAc 10:0 to 60:40, v/v) to afford **12**, a mixture of enantiomers, (20 mg, 13%) as a purple solid. ^1^H NMR (400 MHz, Chloroform-d): d ppm 1.23-1.27 (m, 9H) 1.69-1.74 (dd, *J*=14.67, 4.36 Hz, 1H), 1.81-1.87 (dd, *J*=14.90, 8.71 Hz, 1H), 3.66-3.77 (m, 1H), 5.46 (s, 1H), 5.95 (s, 1H), 6.97 (br d, *J*=7.11 Hz, 1H), 7.17-7.23 (m, 3H), 7.38 (t, *J*=7.07 Hz, 2H), 8.20 (br s, 1H). MS (dual ESI/APCI) *m/z* calculated for C_18_H_22_N_2_O_3_ [M+H]^+^ 315.16, found 315.2.

**Supplementary Text S5.**

Data were additionally exported from mzMine3 into MetaboAnalyst 6.0 for statistical analysis and identification of biomarker metabolites.^4,5^ Missing values for each feature were substituted with 1/5 of the minimum value for each respective variable, and features with greater than 20% missing values were removed. Features that were near-constant throughout experiments, based on an interquartile range of 40%, were filtered out following default parameters for over 1,000 features, and features with low repeatability in QC samples (>20% relative standard deviation) were also removed. Finally, samples were normalized to dry fish weight (mg), transformed using *log* base 10 transformation, and scaled using auto scaling (mean-centered and divided by the standard deviation of each variable).

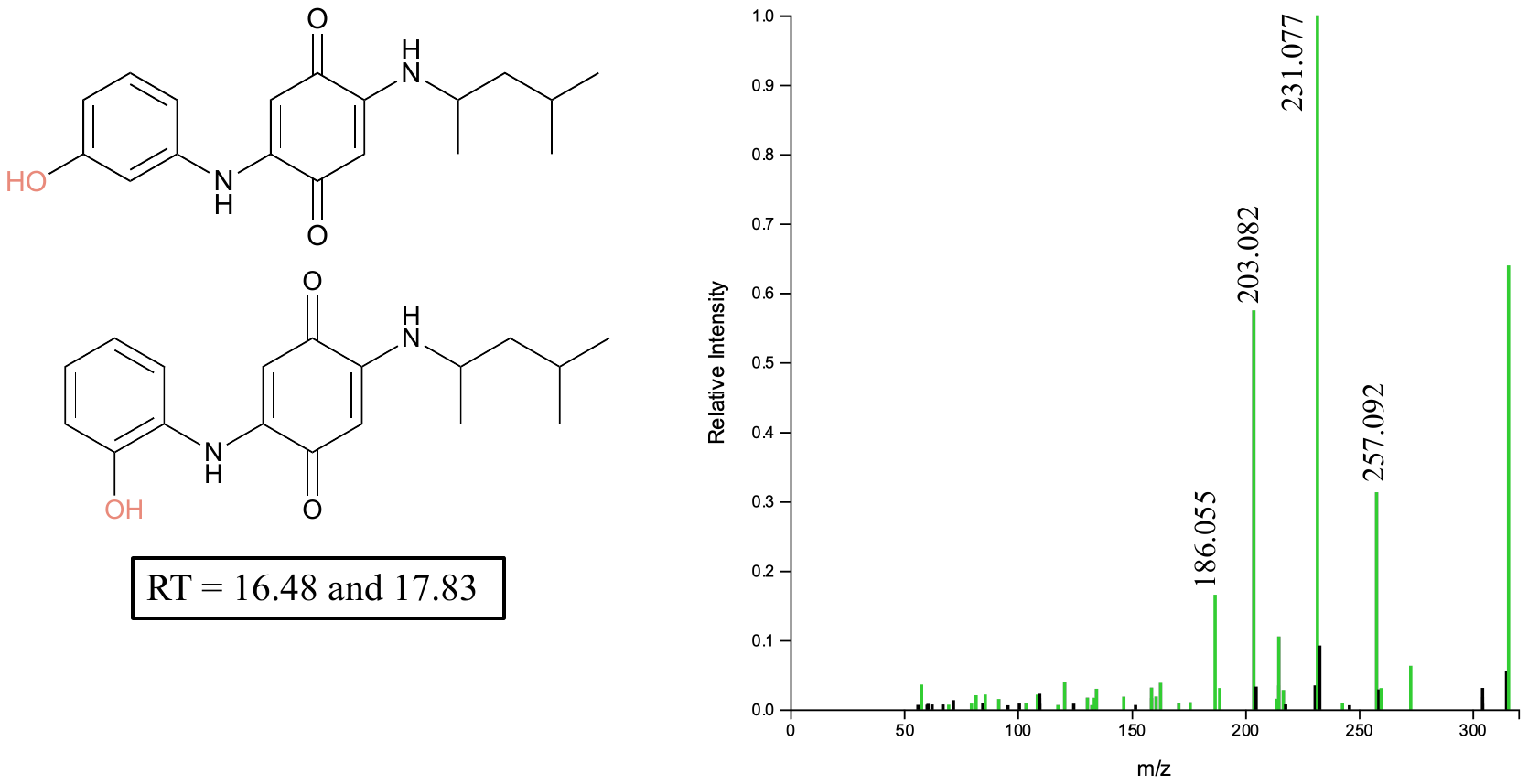

**Supplementary Figure S1.** Phenyl ring monohydroxylation TPs detected in the highest exposure (10 µg/L) in BRT probable structure and corresponding MS^2^. The two additional mono hydroxylation TPs in BRT, when exposed to 10 µg/L, likely indicate that these Phase I hydroxylation’s are not preferred and result from oversaturation of the preferred biotransformation pathway.

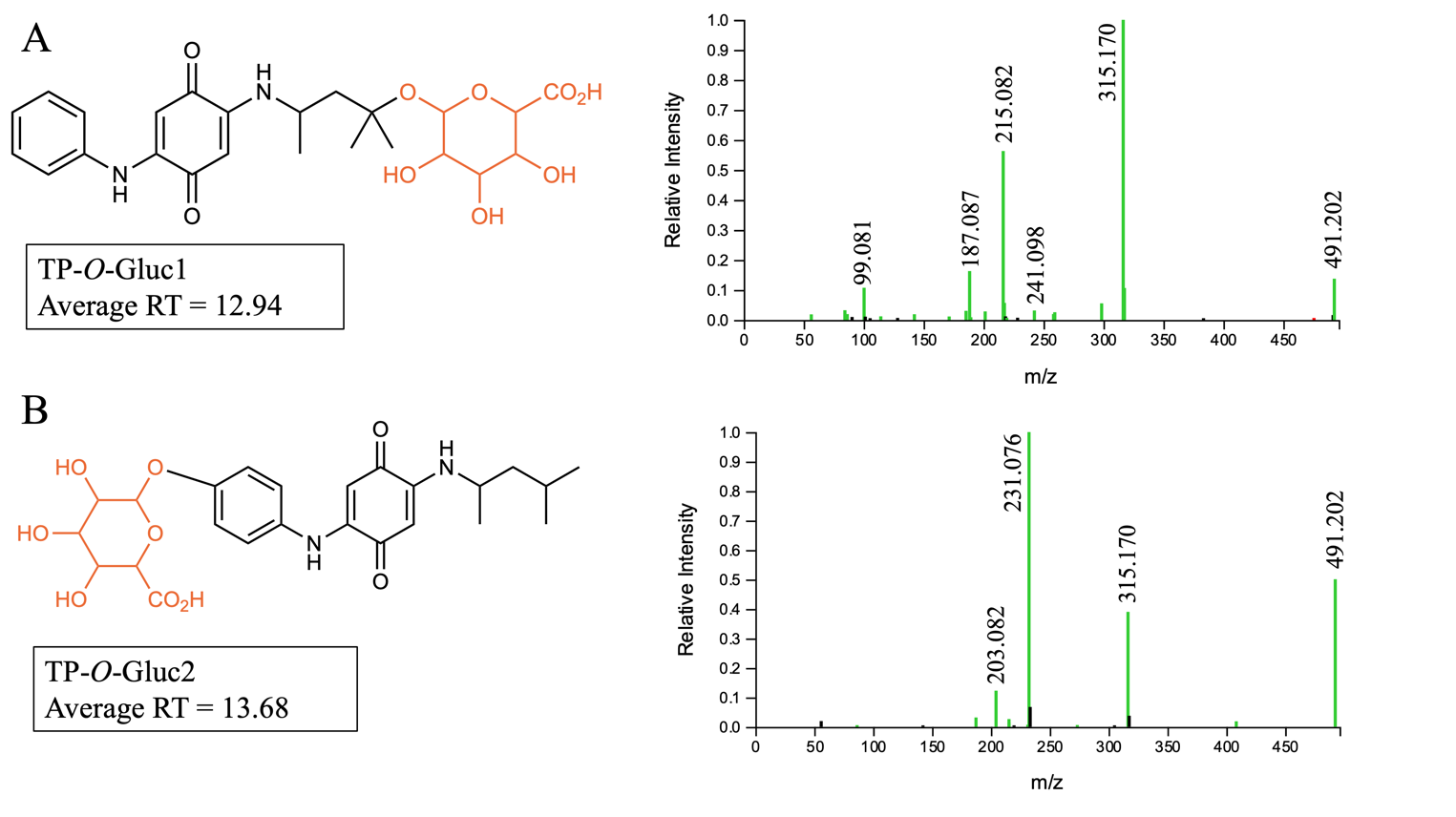

**Supplementary Figure S2.** 6PPD-Q glucuronide-conjugated monohydroxylated TPs (*m/z* = 491.202) with both O-glucuronidation at either the alkyl sidechain hydroxylation (TP-*O*-Gluc1) or phenyl ring hydroxylation (TP-*O*-Gluc2) with neutral loss of the glucuronide acid (*m/z* = 176.032). A) Confirming ions for TP-*O*-Gluc1 from MS2 included *m/z* of 315.170, 241.098, 215.082, and 187.087. B) Confirming ions of TP-*O*-Gluc2 from MS2 included *m/z* of 315.170, 231.076, and 203.082.

**Supplementary Figure S3.** 6PPD-Q glucuronide-conjugated dihydroxylated TPs (*m/z* = 507.198) MS^2^ with likely both *O*-glucuronidation and *N*-glucuronidation TPs being detected.

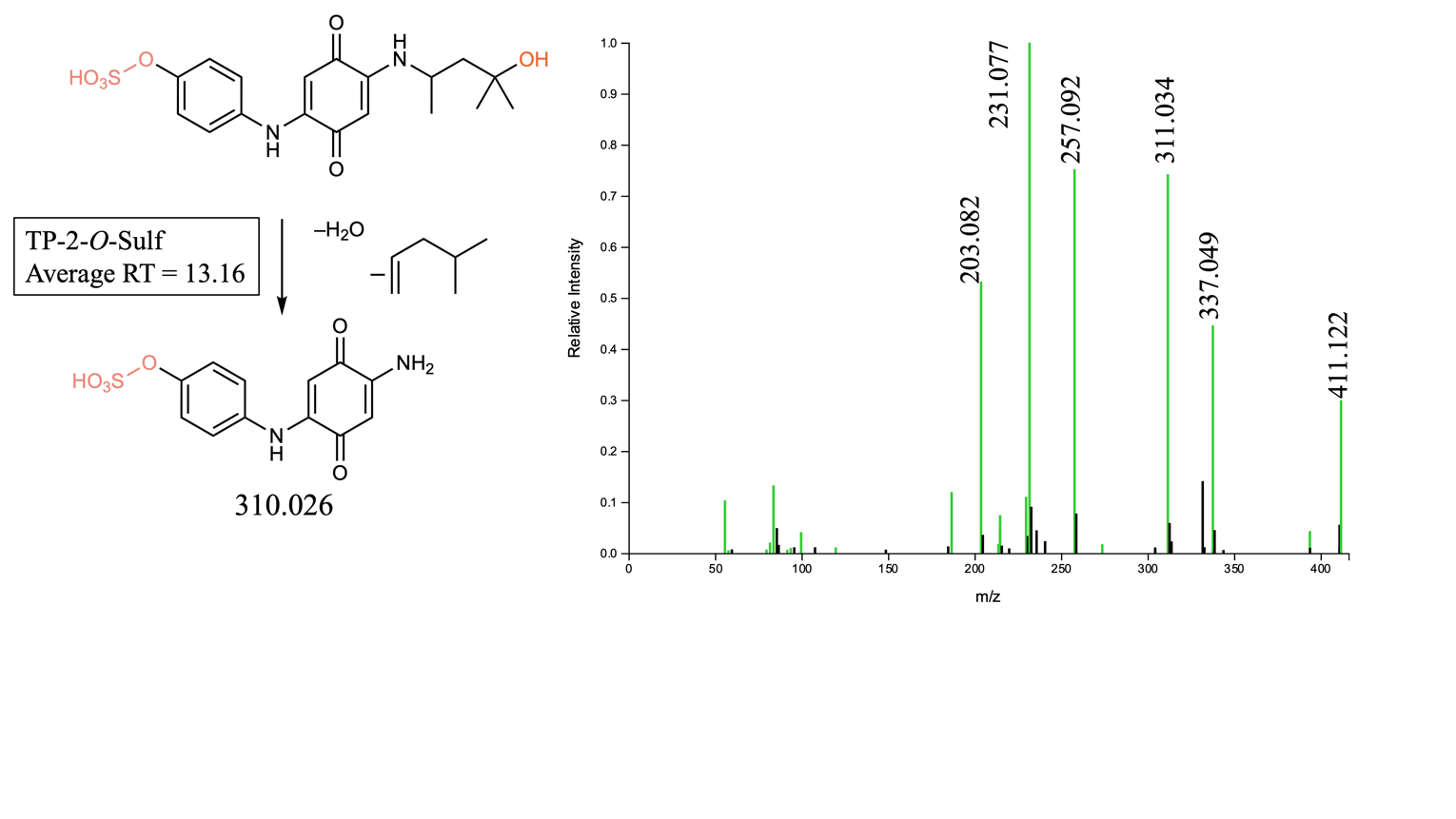

**Supplementary Figure S4.** 6PPD-Q sulfonate-conjugated dihydroxylated TP was *O*-sulfonated at the phenyl ring hydroxylation with an MS^2^ fragment of *m/z* = 311.034 indicating loss of water and alkyl side chain.

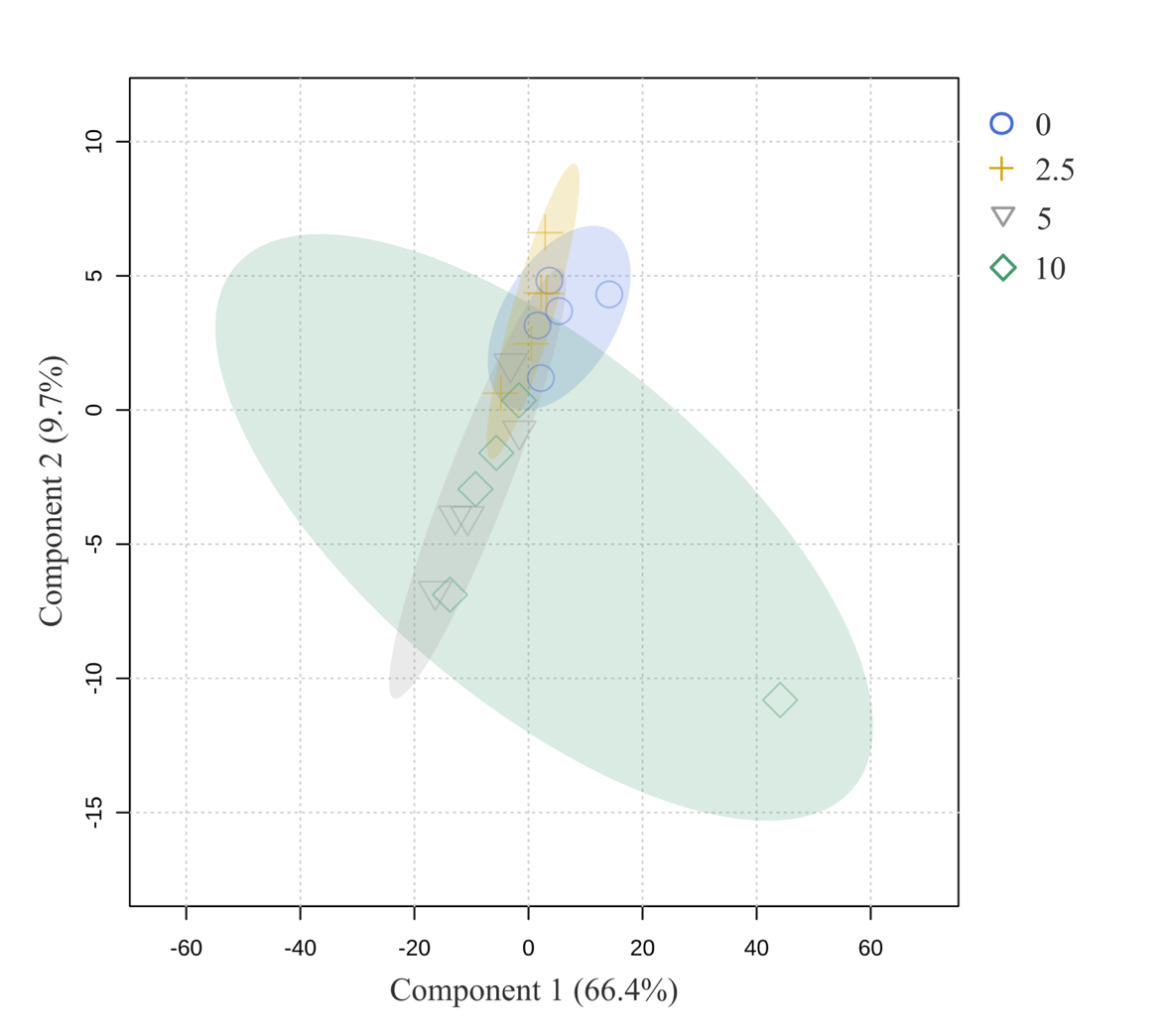

**Supplementary Figure S5.** PLS-DA of BRT metabolome response to 6PPD-Q treatment. Individual points represent each individual sample collected for all treatments. Shapes and shading represent the different treatment groups.

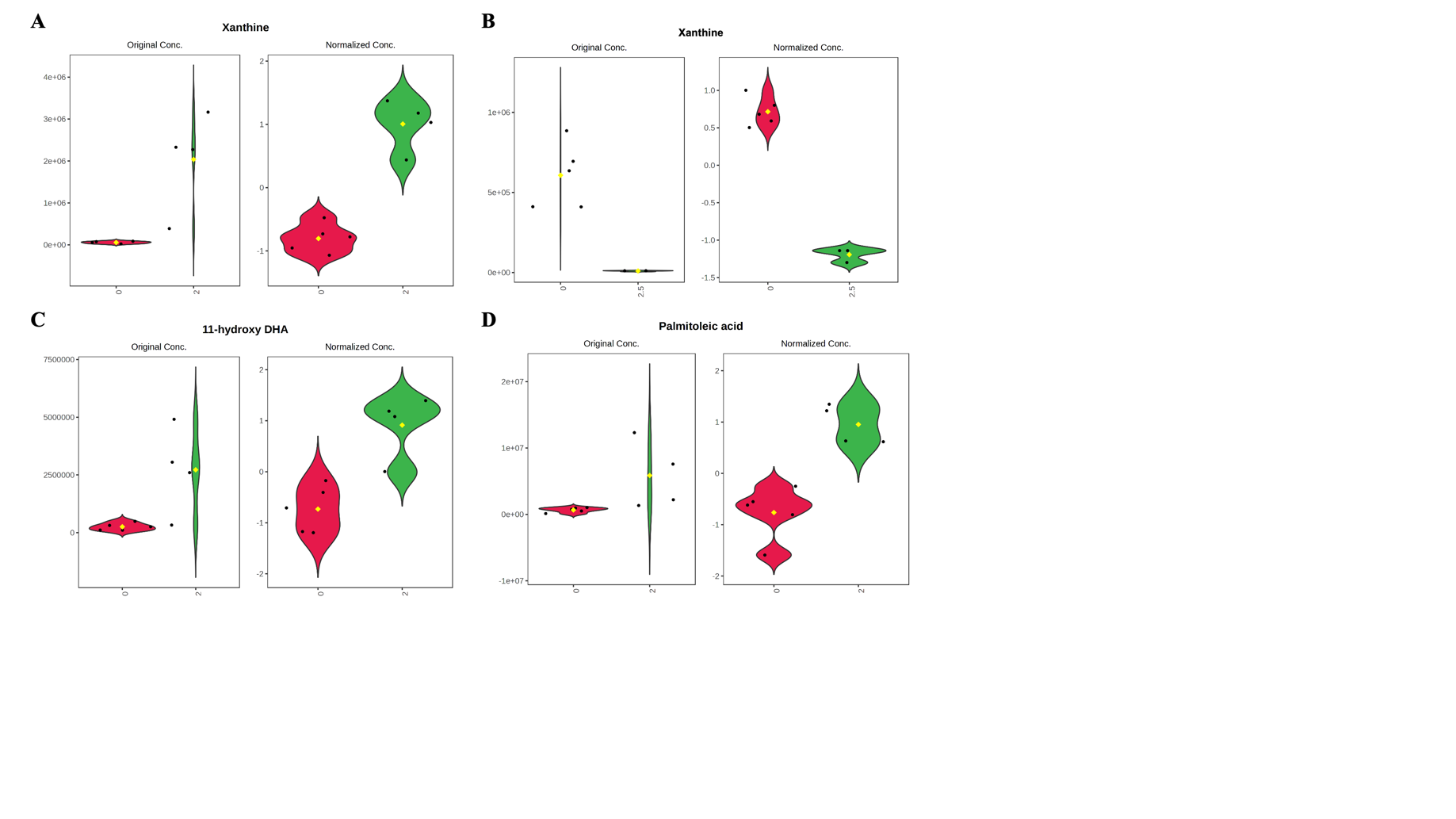

**Supplementary Figure S6.** Violin plots of dysregulated endogenous metabolites in 6PPD-Q exposed fish. A) Xanthine was increased in RBT but B) decreased in 6PPD-Q exposed LKT. C) 11-hydroxy DHA and D) Palmitoleic acid were both increased in 6PPD-Q exposed RBT. The metabolites have the original peak areas and weight normalized, *log* base 10 transformed, and auto-scaled peak areas.

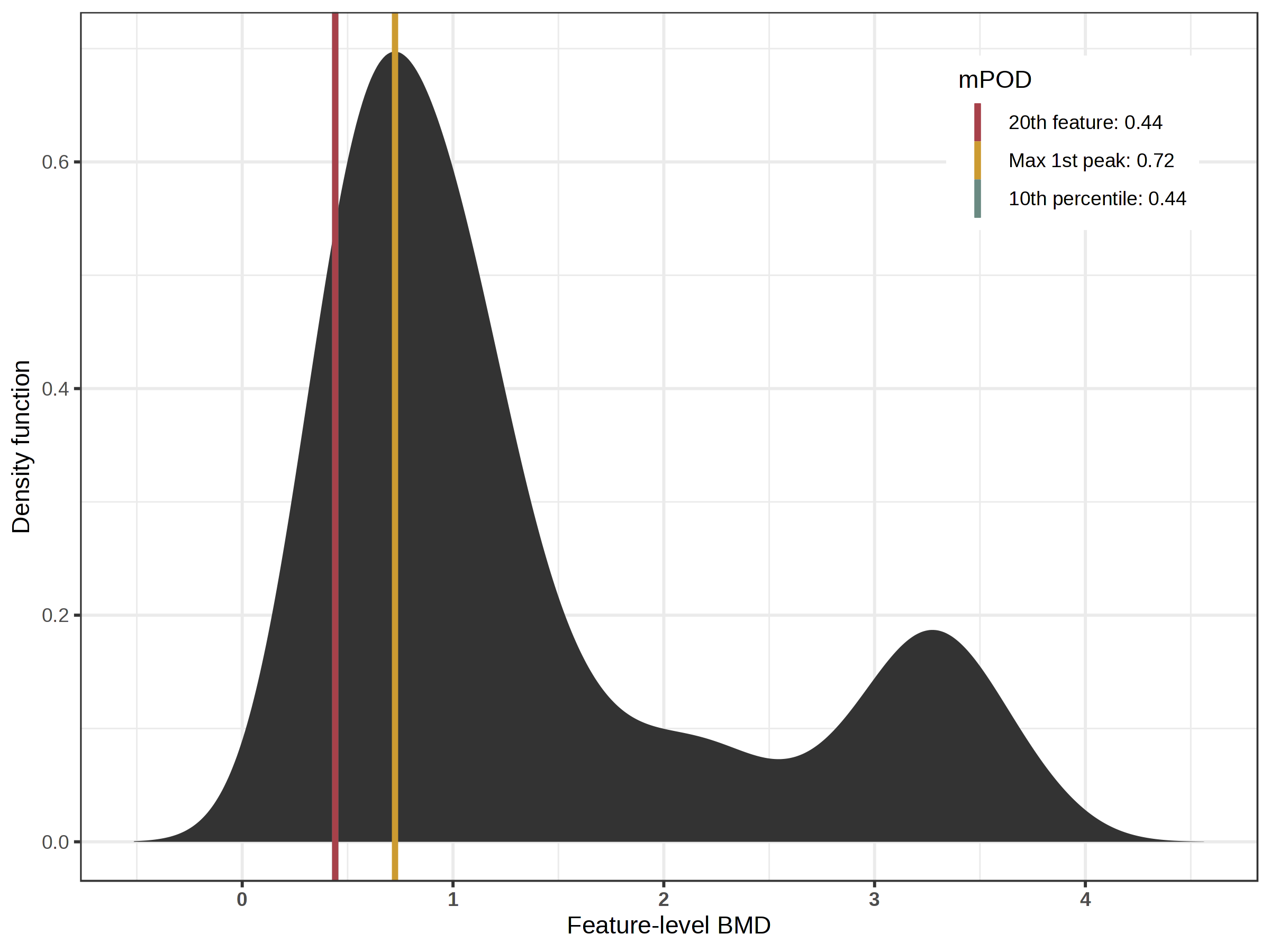

**Supplementary Figure S7.** Metabolomics points-of-departure (mPOD) in 28 d larval RBT exposed to 6PPD-Q. mPODs were estimated from metabolomic BMD analysis as (1) the BMD of the 20^th^ most sensitive metabolite (20^th^ feature; red), (2) the max 1^st^ peak of metabolite BMD distribution (Max 1^st^ peak, yellow), and (3) the 10^th^ percentile of all metabolite BMDs (10^th^ percentile, green).

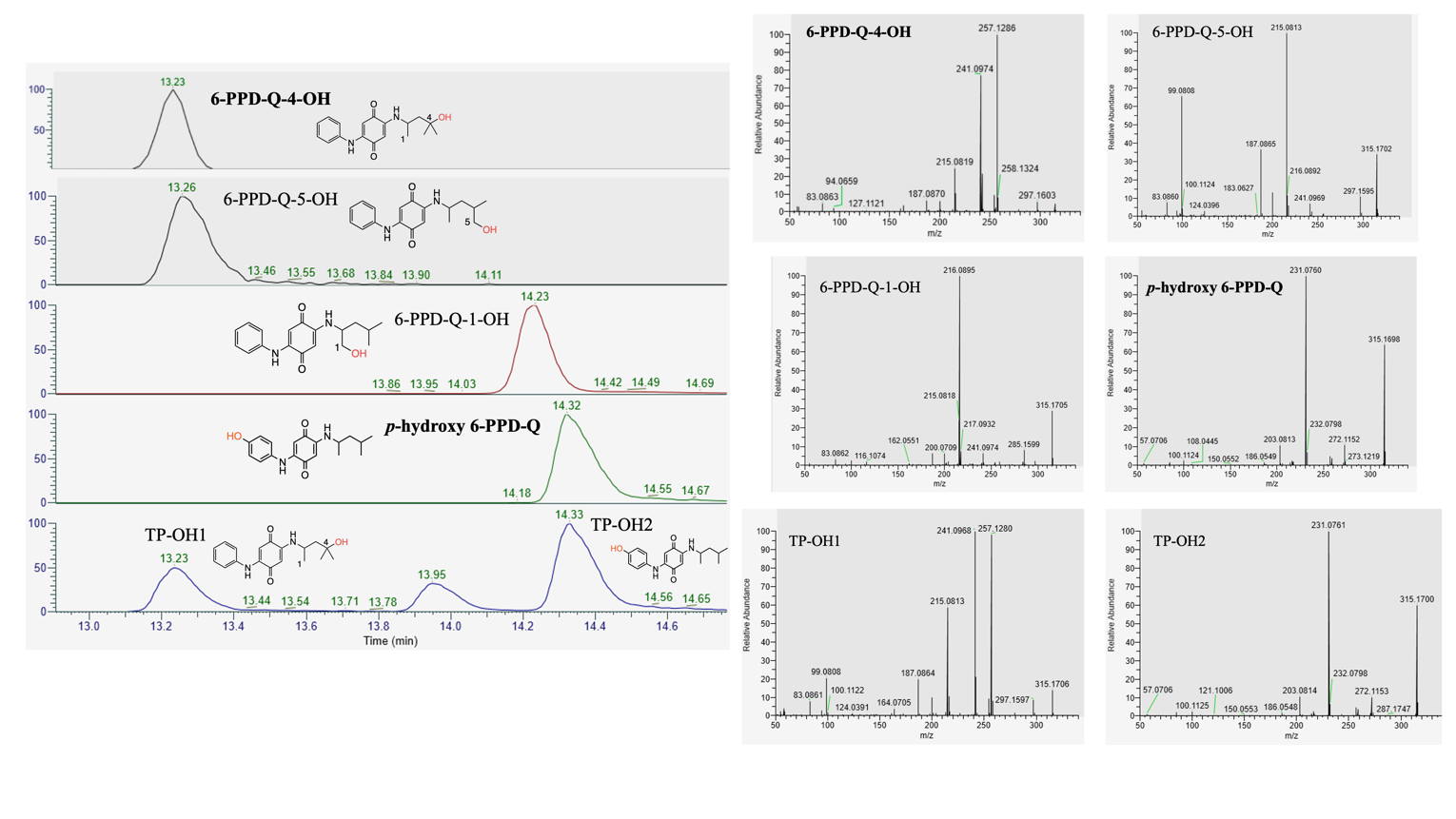

**Supplementary Figure S8.** Retention time and MS^2^ fragmentation of 6-PPD-Q-4-OH (Cayman Chemical), 6-PPD-Q-5-OH (Cayman Chemical), 6PPD-Q-1-OH (Cayman Chemical item number 41591), *p*-hydroxy 6PPD-Q (Cayman Chemical item number 40605), TP-OH1, and TP-OH2. Confirmation of **6-PPD-Q-4-OH** with TP-OH1 with both having a retention time of 13.23 min was observed in the cell media samples with the appropriate fragmentation (*m/z* = 187.087, 200.071, 215.082, 241.097, 257.129 and 297.160) and relative abundance of the fragments. Confirmation of ***p*-hydroxy 6-PPD-Q** with TP-OH2 with a shared retention time of 14.33 min was observed in the cell media samples with the appropriate fragmentation (*m/z* = 203.082, 231.076, and 272.116) and relative abundance of the fragments.

**Supplementary Table S1.** Experimental treatment groups, nominal concentrations of 6PPD-Q exposure for each respective species, and larval fish tissue dry weight (mg; average ± standard deviation).

|  | **Nominal Exposure Concentrations** | | | | | | | | | | |
| --- | --- | --- | --- | --- | --- | --- | --- | --- | --- | --- | --- |
| **Species** | **SC**  **(0 µg/L)** | **0.125** | **0.25** | **0.5** | **1** | **1.25** | **2** | **2.5** | **4** | **5** | **10** |
| **RBT** | **5**  (38.8 ± 7.26) | **5**  (39.6 ± 8.17) | **5**  (39.8 ± 6.18) | **4**  (38.0 ± 5.29) | **5**  (13.0 ± 6.82) |  | **4**  (17.0 ± 9.06) |  | **3**  (31.0 ± 19.3) |  |  |
| **LKT** | **5**  (18.8 ± 7.40) |  |  |  |  | **5**  (40.0 ± 30.0) |  | **3**  (29.0 ± 2.65) |  |  |  |
| **BRT** | **5**  (74.4 ± 8.65) |  |  |  |  |  |  | **5**  (64.2 ± 9.20) |  | **5**  (51.6 ± 9.96) | **4**  (47.3 ± 10.1) |

**Supplementary Table S2.** Water chemistry including temperature, dissolve oxygen, ammonia, nitrite, and nitrate.^6,7^

|  | Temperature (ºC) | Dissolve Oxygen (%) | Ammonia (mg/L) | Nitrite (mg/L) | Nitrate (mg/L) |
| --- | --- | --- | --- | --- | --- |
| Rainbow Trout | 14 ± 0.5 | 90.0 ± 4.6 | 0.08 ± 0.09 | 0.21 ± 0.08 | 0.66 ± 0.25 |
| Lake Trout | 10 ± 0.5 | 93 ± 4.65 | 0.03 ± 0.18 | 0.47 ± 0.58 | 0.73 ± 0.23 |
| Brown Trout | 9.98 ± 0.12 | 100.06 ± 1.19 | 0.6 | 0.071 | 1.2 |

**Supplementary Table S3.** Positive mode targeted gradient elution method for exposure. Flow rate = 0.2 mL/min, column temperature = 40 ^°^C, solvent A = 95% H_2_O: 5% MeOH + 0.1% formic acid and B = 100% MeOH + 0.1% formic acid.

| **Time (min)** | **%B** |
| --- | --- |
| 0.000 | 5 |
| 4.000 | 100 |
| 7.500 | 100 |
| 7.510 | 5 |
| 11.000 | 5 |

**Supplementary Table S4.** Positive mode gradient elution method for non-targeted metabolomics and analytical standard verification. Flow rate = 0.2 mL/min, column temperature = 40 ^°^C, solvent A = 95% H_2_O: 5% MeOH + 0.1% formic acid and B = 100% MeOH + 0.1% formic acid.

| **Time (min)** | **%B** |
| --- | --- |
| 0.00 | 5 |
| 7.50 | 40 |
| 15.00 | 100 |
| 20.00 | 100 |
| 20.10 | 5 |
| 25.00 | 5 |

**Supplementary Table S5.** Precursor and product ions ([M+H]+) and retention time details for 6PPD-Q, 6PPD-Q-d5, Phase I, and Phase II transformation products (TPs) in 2-2.5 µg/L nominal exposed *Salmonidae*, including rainbow trout (*Oncorhynchus mykiss*), lake trout (*Salvelinus namaycush*), and brown trout (*Salmo trutta*).

| **Compound Name (CAS # or Cayman ID)** | **Compound Details** | **Precursor Ion** | **Product Ion** | **Ret Time (min)** |
| --- | --- | --- | --- | --- |
| 6PPD-Q (2754428-18-5) | Environmental TP derived from tire-derived 6PPD reacting with ozone. | 299.175 | 256.121  241.097  215.082  200.071  187.087  100.112 | 15.67 |
| 6PPD-Q-d5 (2750119-14-1) | Deuterium-labeled 6PPD-Q, with all five deuterium atoms located on the phenyl ring. | 304.207 | 261.152  248.144  246.129  220.113  205.102  192.118 | 15.65 |
| TP-OH1 or 6PPD-Q-4-OH | Alkyl sidechain monohydroxylation at carbon 4. Primarily metabolized via CYP1A enzyme. | 315.170 | 257.129  241.097  215.082  200.071  187.087  164.071  99.081 | 13.45 |
| TP-OH2 or  *p*-hydroxy 6PPD-Q (40605) | Phenyl ring monohydroxylation at the para position. This is the more dominant isomer. | 315.170 | 272.116  257.092  231.077  203.082 | 14.56 |
| TP-2-OH | An alkyl side chain and phenyl ring dihydroxylation TP being a combination of both TP-OH1 and TP-OH2. | 331.165 | 313.154  273.123  257.092  243.077  231.077  216.066  203.082 | 12.11 |
| TP-*O*-Gluc1 | TP-OH1 glucuronidation TP. | 491.202 | 315.170  241.098  215.082  187.087  99.081 | 12.94 |
| TP-*O*-Gluc2 | TP-OH2 glucuronidation TP. This is the more dominant isomer. | 491.202 | 315.170  231.076  203.082 | 13.68 |
| TP-2-*O*-Gluc1 | Dihydroxylated (TP-2-OH) glucuronidation. | 507.198 | 331.165  257.092  231.077  203.082 | 10.42 |
| TP-2-*O*-Gluc2 | Dihydroxylated (TP-2-OH) glucuronidation. | 507.198 | 331.165  231.077  203.082 | 11.42 |
| TP-2-*O*-Gluc3 | Dihydroxylated (TP-2-OH) glucuronidation. | 507.198 | 331.165  241.098  231.077  215.082  203.081 | 11.80 |
| TP-2-*O*-Gluc4 | Dihydroxylated (TP-2-OH) glucuronidation. | 507.198 | 331.166  247.072 | 13.86 |
| TP-2-*O*-Sulf | Dihydroxylated (TP-2-OH) sulfation. Likely conjugated on hydroxyl group at para position of phenyl ring. | 411.122 | 337.049  311.034  257.092  231.077  203.082 | 13.16 |
